# The neuronal architecture of autonomic dysreflexia

**DOI:** 10.1101/2024.05.06.592781

**Authors:** Jan Elaine Soriano, Rémi Hudelle, Loïs Mahe, Matthieu Gautier, Yue Yang Teo, Michael A. Skinnider, Achilleas Laskaratos, Steven Ceto, Claudia Kathe, Thomas Hutson, Rebecca Charbonneau, Fady Girgis, Steve Casha, Julien Rimok, Marcus Tso, Kelly Larkin-Kaiser, Nicolas Hankov, Aasta P. Gandhi, Suje Amir, Xiaoyang Kang, Yashwanth Vyza, Eduardo Martin-Moraud, Stephanie Lacour, Robin Demesmaeker, Leonie Asboth, Quentin Barraud, Mark A. Anderson, Jocelyne Bloch, Jordan W. Squair, Aaron A. Phillips, Grégoire Courtine

**Author notes:** These authors contributed equally (Jan Elaine Soriano, Rémi Hudelle). These authors jointly supervised this work (Jordan W. Squair, Aaron A. Phillips, Grégoire Courtine).

## Abstract

Autonomic dysreflexia is a life-threatening medical condition characterized by episodes of uncontrolled hypertension that occur in response to sensory stimuli after spinal cord injury (SCI)^1–7^. The fragmented understanding of the mechanisms underlying autonomic dysreflexia hampers the development of therapeutic strategies to manage this condition, leaving people with SCI at daily risk of heart attack and stroke^8–18^. Here, we expose the complete *de novo* neuronal architecture that develops after SCI and causes autonomic dysreflexia. In parallel, we uncover a competing, yet overlapping neuronal architecture activated by epidural electrical stimulation of the spinal cord that safely regulates blood pressure after SCI. The discovery that these adversarial neuronal architectures converge onto a single neuronal subpopulation provided a blueprint for the design of a mechanism-based intervention that reversed autonomic dysreflexia in mice, rats, and humans with SCI. These results establish a path for the effective treatment of autonomic dysreflexia in people with SCI.

---

Spinal cord injury (SCI) disrupts the communication between the brainstem vasomotor centers and the regions of the spinal cord that regulate hemodynamics^19^. The resulting isolation of neurons in the spinal cord triggers a progressive maladaptive reorganization of neuronal projections throughout the spinal cord below the injury that permits the insidious emergence of uncontrolled hypertensive episodes, known as autonomic dysreflexia^1–7^. The resulting isolation of neurons in the spinal cord triggers a progressive maladaptive reorganization of neuronal projections throughout the spinal cord below the injury that permits the insidious emergence of uncontrolled hypertensive episodes, known as autonomic dysreflexia^1–7^. The consequence of these hypertensive episodes is a daily risk of life-threatening cardiovascular events^8–18^.

We reasoned that disentangling the specific neuronal sub-populations involved in the emergence of autonomic dysreflexia, and how these neurons and their projection patterns reorganize after SCI, would uncover key principles to target these neurons therapeutically and thus eliminate autonomic dysreflexia due to SCI.

## Spatial organization of neurons involved in autonomic dysreflexia

In humans with SCI, episodes of autonomic dysreflexia are most commonly triggered by bladder or bowel distension, lower urinary tract infections, and pressure sores^2^. We reasoned that identifying the neurons triggering autonomic dysreflexia would require a preclinical model that provokes reliable, repeatable, and predictable episodes of autonomic dysreflexia.

To establish this model, we elicited autonomic dysreflexia using colorectal distension in mice^20,21^with complete upper-thoracic SCI and monitored pressor responses with beat-by-beat blood pressure monitoring (**Fig. 1a-b and Supplementary Fig. 1a-b**). We found that autonomic dysreflexia emerged approximately two weeks after SCI (**Fig. 1c and Supplementary Fig. 1c**), after which the amplitude of these pressor responses increased gradually until reaching a plateau by one month after SCI (**Fig. 1c and Supplementary Fig. 1c**).

**Fig. 1.**
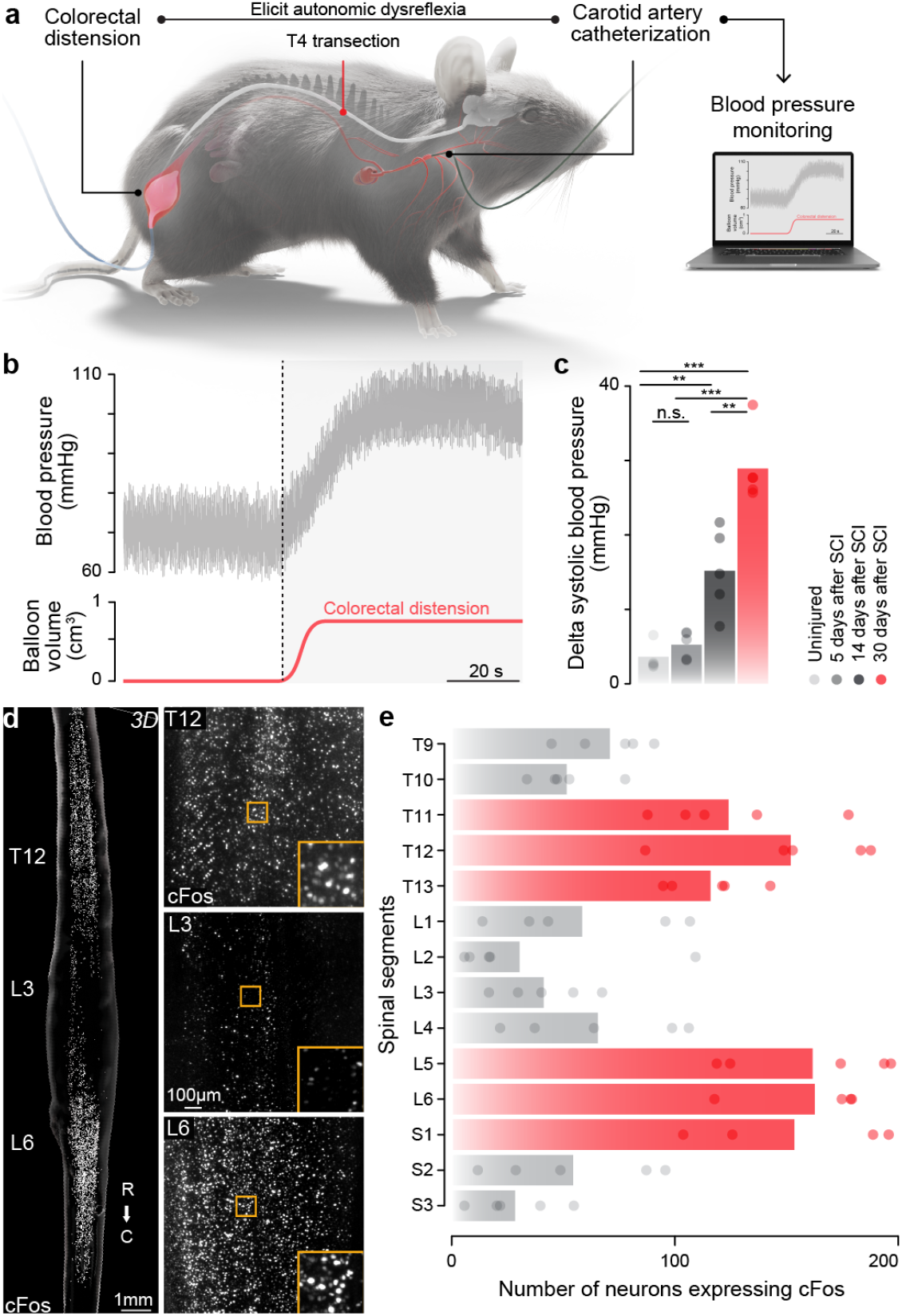
Autonomic dysreflexia triggers transcriptional activity in the lumbosacral and lower thoracic spinal cord. **a**, Experimental model to quantify the severity of autonomic dysreflexia in mice with complete SCI. **b**, Changes in blood pressure during an episode of autonomic dysreflexia elicited by a controlled colorectal distension. **c**, Severity of autonomic dysreflexia measured by the change in systolic blood pressure elicited by a controlled colorectal distension at different timepoints after SCI. Raw data and statistics provided in **Supplementary Table 1**. **d**, Whole spinal cord visualization of immunohistochemical staining for cFos^22,23^ in a mouse with SCI that was exposed to repetitive episodes of autonomic dysreflexia. **e**, Bar plots reporting the mean number of cFos-labelled neurons for each spinal cord segment quantified in mice with SCI that were exposed to repetitive episodes of autonomic dysreflexia (n = 5; mixed effect linear model; p-value < 0.001), demonstrating a clear enrichment in the lumbosacral and lower thoracic spinal cord.

To identify the regions of the spinal cord activated during autonomic dysreflexia, we optimized immunolabeling-enabled three dimensional imaging of solvent-cleared organs (iDISCO+)^22,23^ to achieve whole-spinal cord labeling of Fos—a marker of neuronal activity-induced transcription^24,25^—and combined this whole spinal cord visualization with high-resolution CLARITY-optimized light-sheet microscopy^26^ (**Fig. 1d and Supplementary Fig. 1d**) and automated 3D nuclear spot detection and quantification^27^ (**Supplementary Fig. 1e**). We quantified activity-induced transcription in the spinal cords of mice with SCI that underwent repetitive autonomic dysreflexia over 90 minutes (**Fig. 1e, Supplementary Fig. 1e**)^3^.

We found that autonomic dysreflexia triggered massive transcriptional activation throughout the spinal cord (**Supplementary Fig. 1e**). However, we detected disproportionate enrichments of neuronal activity in two well-defined regions. The first enrichment occurred in the lumbosacral segments that receive the sensory afferents conveying information from the colorectal stimulus (**Fig. 1d**). The second enrichment emerged within the lower thoracic spinal cord, which is referred to as the *hemodynamic hotspot*^19^. This region hosts a dense concentration of sympathetic preganglionic neurons that access the splanchnic vasculature through ganglionic neurons to elicit powerful pressor responses^19^. We confirmed these segment-specific enrichments of neuronal activity with classical immunofluorescence of Fos on sectioned tissue (**Supplementary Fig. 1f**).

The presence of discrete yet distant enrichments in neuronal activity demonstrates that autonomic dysreflexia due to bowel distension activates neuronal subpopulations in the lumbosacral spinal cord, and suggests that these neurons establish axonal projections onto neurons located within the lower thoracic spinal cord, coinciding with the *hemodynamic hotspot* that regulates blood pressure.

## The neurons activated by autonomic dysreflexia

We anticipated that understanding the emergence of maladaptive communication between the lumbosacral and lower thoracic spinal cord would be contingent on identifying the neuronal subpopulations in each region that are activated during autonomic dysreflexia.

The neuronal subpopulations embedded within the spinal cord are parcellated into a hierarchical organization that arises from their neurotransmitter expression, developmental transcription factors, and projection patterns^28–30^. This hierarchical organization dictates that identifying the neurons involved in specific neurological functions must follow the logical progression along the cardinal classes.

To follow this progression, we first asked whether the neurons embedded in the lumbosacral and lower thoracic spinal cords, and activated during autonomic dysreflexia, exhibited an excitatory or inhibitory phenotype.

To answer this question, we exposed Vglut2^Cre^::Ai9^(RCL-tdT)^ and Vgat^Cre^::Ai9^(RCL-tdT)^ repetitive autonomic dysreflexia. We quantified the proportion of activated neurons that colocalized with each neurotransmitter phenotype in the lumbosacral and lower thoracic spinal cords (**Supplementary Fig. 2a-b**). We found that autonomic dysreflexia triggered a disproportionate activation of Vglut2^ON^ neurons in the lumbosacral and lower thoracic spinal cords compared to Vgat^ON^ (Supplementary Fig. 2b)^3,7^.

We next sought to determine the causal role of these neuronal subpopulations in triggering autonomic dysreflexia. To manipulate each neuronal subpopulation in each location, we expressed inhibitory Designer Receptors Exclusively Activated by Designer Drugs (DREADDs)^31^ in Vglut2^ON^, Vgat^ON^ or Chat^ON^ neurons located either in the lumbosacral or lower thoracic spinal cord using targeted infusions of AAV5-hSyn-DIO-hM4Di-mCherry in Cre-driver mouse lines **(Supplementary Fig. 2c)**^32^.

Chemogenetic inactivation restricted to Vglut2^ON^, neurons located in the lumbosacral or lower thoracic spinal cord blunted autonomic dysreflexia (**Supplementary Fig. 2c**). In contrast, chemogenetic silencing of Vgat^ON^, neurons had no effect (**Supplementary Fig. 2d**).

These experiments suggested that Vglut2^ON^ neurons located in the lumbosacral spinal cord must establish axonal projections within the lower thoracic spinal cord.

To expose these putative projections, we labeled the axons and synapses of Vglut2^ON^ neurons with infusions of AAV-DJ-hSyn-flex-mGFP-2A-synaptophysin-mRuby^33^ into the lumbosacral spinal cord of Vglut2^Cre^ mice **(Supplementary Fig. 2d)**. We detected long-distance projecting axons that established synapses throughout the thoracolumbar spinal cord. SCI altered this projection pattern, whereby the density of synaptic terminals increased significantly within the lower thoracic spinal cord segments (**Supplementary Fig. 2d**).

Our results thus far revealed that Vglut2^ON^ neurons located in the lumbosacral spinal cord propel axonal projections to the lower thoracic spinal cord where Vglut2^ON^ neurons are activated to trigger autonomic dysreflexia.

However, Vglut2^ON^ neurons comprise diverse neuronal subpopulations that collectively encompass more than 50% of neurons in the spinal cord^28–30^. Consequently, we felt compelled to descend the hierarchical organization of neurons in the spinal cord in order to identify the precise neuronal subpopulations that govern the emergence of autonomic dysreflexia. However, this descent is contingent on an atlas that catalogs the molecular perturbation elicited by autonomic dysreflexia across the compendium of neuronal subpopulations in the spinal cord.

To establish this comparative atlas, we profiled the lumbosacral and lower thoracic spinal cord of mice exposed to repetitive episodes of autonomic dysreflexia using single-nucleus RNA sequencing (snRNA-seq)^34–38^. We obtained high-quality transcriptomes from 64,739 nuclei that were evenly represented across experimental conditions and spatial locations (**Supplementary Fig. 3a-b**). We identified all of the major cell types of the mouse spinal cord (**Supplementary Fig. 3c-j**). We then integrated this dataset within our previous atlases of the mouse spinal cord^27,29,30,39,40^, which enabled us to annotate highly specific neuronal subpopulations that parcellated into dorsal versus ventral, excitatory versus inhibitory, and local (*Nfib*) versus long-projecting (*Zfhx3*) populations (**Fig. 2a, Supplementary Fig. 3k-l**).

**Fig. 2.**
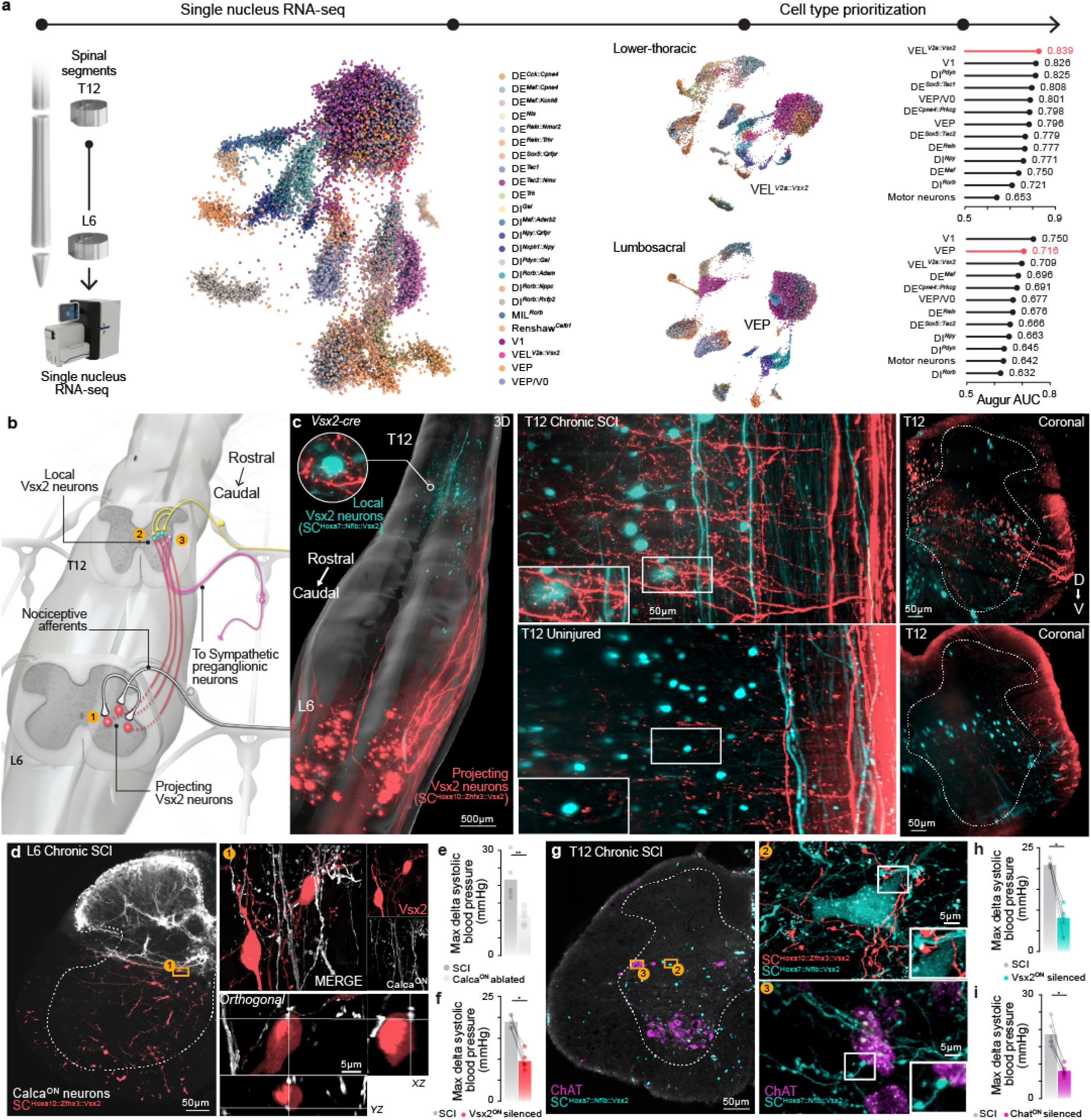
The neuronal architecture of autonomic dysreflexia. **a**, Schematic overview of the single-nucleus sequencing experiment. Uniform manifold approximation and projection (UMAP) visualization of 64,739 neuronal nuclei, colored by neuronal subpopulation identity. *Middle*, UMAP visualizations of neuronal subpopulations in the lower thoracic (top) and lumbosacral (bottom) spinal cord. *Right*, Ranking neuronal subpopulations most responsive to autonomic dysreflexia with Augur. **b**, Schematic overview of the neuronal architecture of autonomic dysreflexia, including the nodes (numbers) that are dissected anatomically and functionally in the subsequent panels. **c**, Whole spinal cord visualization of projections from SC^Hoxa10::Zfhx3::Vsx2^ neurons located in the lumbosacral spinal cord that project to SC^Hoxa7::Nfib::Vsx2^ neurons located in the lower thoracic spinal cord. Insets illustrate the synaptic-like appositions of Calca^ON^ projections labeled with immunohistochemistry onto SC^Hoxa10::Zfhx3::Vsx2^ neurons in the lumbosacral spinal cord. **d**, Calca^ON^ projections labeled with immunohistochemistry onto SC^Hoxa10::Zfhx3::Vsx2^ neurons in the lumbosacral spinal cord, including insets showing synaptic-like appositions. **e**, Bar plot reporting the severity of autonomic dysreflexia, quantified as the mean change in systolic blood pressure in response to colorectal distension before and after the ablation of Calca^ON^ neurons located in the dorsal root ganglia in Calca^Cre^::Advil^FlpO^::iDTR mice (n = 5; independent samples t-test; t = -6.0; p-value = 0.0006). **f**, Severity of autonomic dysreflexia before and after chemogenetic silencing of Vsx2^ON^ neurons located in the lumbosacral spinal cord in Vsx2^Cre^ (n = 5; paired samples t-test; t = -9.47; p-value = 0.00069). **g**, Projections from SC^Hoxa10::Zfhx3::Vsx2^ in the lower thoracic spinal cord co-labeled with SC^Hoxa7::Nfib::Vsx2^ neurons and their local projections as well as immunohistochemical labelling of Chat^ON^. Insets show synaptic-like appositions from SC^Hoxa10::Zfhx3::Vsx2^ neurons onto SC^Hoxa7::Nfib::Vsx2^, and synaptic-like appositions of projections from SC^Hoxa7::Nfib::Vsx2^ to Chat^ON^ sympathetic preganglionic neurons located in the intermediolateral column. **h**, Severity of autonomic dysreflexia before and after chemogenetic silencing of Vsx2^ON^ neurons located in the lower thoracic spinal cord in Vsx2^Cre^ (n = 5; paired samples t-test; t = -9.39; p-value = 0.00072). **i**, Severity of autonomic dysreflexia before and after chemogenetic silencing of Chat^ON^ neurons located in the lower thoracic spinal cord in Chat^Cre^ mice (n = 5; paired samples t-test; t = -8.03; p-value = 0.00048).

To identify the neuronal subpopulations perturbed by autonomic dysreflexia, we applied cell type prioritization^39,41^.

We captured the principle of cell type prioritization in a machine-learning method called Augur, which identifies cell types undergoing transcriptional responses to a perturbation by ranking cell types that are increasingly more separable within the highly multidimensional space of gene expression. This prioritization exposed Vsx2^ON^ excitatory neurons as the most transcriptionally responsive neuronal sub-population during autonomic dysreflexia (**Fig. 2a**). Although Vsx2^ON^ neuronal subpopulations were prioritized both in the lumbosacral (neurons defined by the expression of Hoxa10) and lower thoracic (Hoxa7) spinal cords, the prioritized neurons in the lumbosacral region were consistent with long-projecting Vsx2^ON^ neurons, since they expressed the marker *Zfhx3* (SC^Hoxa10::Zfhx3::Vsx2^), while the most perturbed neuronal subpopulation in the lower thoracic spinal cord instead expressed markers of locally-projecting neurons *Nfib* (SC^Hoxa7::Nfib::Vsx2^) (**Fig. 2a, Supplementary Fig. 3m-n**)^28^. Visualization of the activity-dependent marker cFos confirmed the activation of SC^Hoxa10::Zfhx3::Vsx2^ and SC^Hoxa7::Nfib::Vsx2^ neurons in the lumbosacral and lower thoracic spinal cord, respectively, in response to autonomic dysreflexia (**Supplementary Fig. 3o**).

Although developmentally defined V2a neurons that express Vsx2 (previously known as Chx10)^42–45^ have been implicated in the production of reaching^46–48^ and walking^46,49–59^, Vsx2^ON^ neuronal subpopulations have not been demonstrated to participate in the regulation of blood pressure. Nonetheless, the results of our comparative snRNA-seq experiments dictate that the activation of SC^Hoxa7::Nfib::Vsx2^ neurons and SC^Hoxa10::Zfhx3::Vsx2^ neurons triggers autonomic dysreflexia.

## The neuronal architecture of autonomic dysreflexia

Since autonomic dysreflexia only emerges after SCI, we reasoned that the injury must induce the formation of a maladaptive neuronal architecture that incorporates SC^Hoxa7::Nfib::Vsx2^ neurons and SC^Hoxa10::Zfhx3::Vsx2^ neurons, and possesses the anatomical and functional features compatible with the requirements to trigger autonomic dysreflexia. To expose this architecture, we conducted sequential anatomical and functional experiments that aimed to reconstruct the successive nodes composing the blueprint of the neuronal architecture responsible for autonomic dysreflexia (**Fig. 2b-c, Supplementary Fig. 4a**).

Small diameter nociceptive afferents act as the primary source of sensory input responsible for triggering autonomic dysreflexia^5,16,60,61^. Consequently, nociceptive neurons are positioned as the first node within the neuronal architecture of autonomic dysreflexia, implying that SC^Hoxa10::Zfhx3::Vsx2^ neurons are likely to receive synaptic projections from these afferents. To expose this connectome, we labeled synapses from Calca^ON^ axonal projections. While SC^Hoxa10::Zfhx3::Vsx2^ neurons in the lumbosacral spinal cord did not receive any Calca^ON^ axonal projections in uninjured mice, we found that SCI triggered the invasion of Calca^ON^ axons into intermediate laminae, where they established *de novo* synaptic appositions onto SC^Hoxa10::Zfhx3::Vsx2^ neurons (**Fig. 2c, Supplementary Fig. 4b-d**). This anatomical reorganization suggested that Calca^ON^ neurons trigger autonomic dysreflexia through the activation of SC^Hoxa10::Zfhx3::Vsx2^ neurons. To expose this causality, we ablated Calca^ON^ neurons in the dorsal root ganglia with diphtheria toxin in Calca^Cre^::Advil^FlpO^::iDTR mice^62^. The ablation of this afferent subpopulation abolished autonomic dysreflexia. In contrast, the ablation of parvalbumin (PV^ON^) neurons in the dorsal root ganglia, which convey proprioceptive information along large diameter afferents that project into the spinal cord, failed to influence autonomic dysreflexia (**Fig. 2d, Supplementary Fig. 4e-f**).

We next posited that SC^Hoxa10::Zfhx3::Vsx2^ act as the second node in the neuronal architecture of autonomic dysreflexia (**Fig. 2b-c, Supplementary Fig. 4g-h**). To expose the necessity of SC^Hoxa10::Zfhx3::Vsx2^ neurons in triggering autonomic dysreflexia, we infused AAV5-hSyn-DIO-hM4D(Gi)-mCherry^63^ into the lumbosacral spinal cord of Vsx2^Cre^ mice. Inactivation of SC^Hoxa10::Zfhx3::Vsx2^ neurons blunted the severity of autonomic dysreflexia (**Fig. 2e, Supplementary Fig. 4i-k**).

We then sought to establish the sufficiency of SC^Hoxa10::Zfhx3::Vsx2^ neurons to trigger autonomic dysreflexia. To expose this sufficiency, we infused AAV5-hSyn-flex-Chrimson-tdTomato^64^ into the lumbar spinal cord of Vsx2^Cre^ mice to express excitatory opsins in SC^Hoxa10::Zfhx3::Vsx2^ neurons. Optogenetic stimulation of SC^Hoxa10::Zfhx3::Vsx2^ neurons immediately triggered autonomic dysreflexia. This increase in blood pressure contrasted with the absence of pressor responses when the same optogenetic manipulation of SC^Hoxa10::Zfhx3::Vsx2^ neurons was performed in uninjured mice (**Supplementary Fig. 4l-n**).

The necessity and sufficiency of SC^Hoxa10::Zfhx3::Vsx2^ neurons to trigger autonomic dysreflexia implied that SCI must also provoke SC^Hoxa10::Zfhx3::Vsx2^ neurons to propel axonal projections to the lower thoracic spinal cord. To uncover these putative projections, we infused rAAV2-flex-hSyn-FlpO into the lower thoracic spinal cord followed by infusions of AAV5-hSyn-Con/Fon-eYFP (enhanced yellow fluorescent protein)^65^ into the lumbosacral spinal cord of Vsx2^Cre^ mice. This neuroanatomical tracing strategy exposed *de novo* long-distance projections from SC^Hoxa10::Zfhx3::Vsx2^ neurons that established synaptic-like appositions onto SC^Hoxa7::Nfib::Vsx2^ neurons in the lower thoracic spinal cord (**Fig. 2c, Supplementary Fig. 4i-k**).

Since SC^Hoxa7::Nfib::Vsx2^ neurons in the lower thoracic spinal cord were transcriptionally perturbed by autonomic dysreflexia and received direct synaptic projections from SC^Hoxa10::Zfhx3::Vsx2^ neurons, we hypothesized that SC^Hoxa7::Nfib::Vsx2^ neurons must act as the third node in the neuronal architecture of autonomic dysreflexia (**Fig. 2c, Supplementary Fig. 5a-e**). To establish the necessity and sufficiency of this node, we manipulated the activity of SC^Hoxa7::Nfib::Vsx2^ neurons. Chemogenetic inactivation of these neurons blunted the severity of autonomic dysreflexia (**Fig. 2g, Supplementary Fig. 5f-h**). In turn, optogenetic activation of SC^Hoxa7::Nfib::Vsx2^ neurons immediately triggered pressor responses (**Supplementary Fig. 5i-k**).

We next asked whether the projection pattern of SC^Hoxa7::Nfib::Vsx2^ neurons in the lower thoracic spinal cord would be compatible with an involvement in autonomic dysreflexia. To answer this question, we infused AAV5-hSyn-flex-tdTomato into Vsx2^Cre^ mice and quantified the density of projections that formed synaptic-like appositions with Chat^ON^ neurons in the intermediolateral column of lower thoracic spinal segments, where Chat^ON^ sympathetic preganglionic neurons reside (**Supplementary Fig. 5l-o**). SC^Hoxa7::Nfib::Vsx2^ neurons primarily projected ventrally, but also expanded projections laterally where they established synaptic-like appositions with the majority of Chat^ON^ neurons located in the intermediolateral column of the lower thoracic spinal cord.

Chat^ON^ sympathetic preganglionic neurons are embedded in the intermediolateral column, and we found that they receive projections from SC^Hoxa7::Nfib::Vsx2^ neurons. Consequently, we surmised that Chat^ON^ neurons act as the fourth and final node in the neuronal architecture of autonomic dysreflexia. Indeed, chemogenetic inactivation of Chat^ON^ neurons located in the lower thoracic spinal cord abolished autonomic dysreflexia elicited by bowel distension (**Fig. 2h, Supplementary Fig. 5p-r**).

These sequential anatomical and functional experiments exposed the complete neuronal architecture that causes autonomic dysreflexia. The building blocks of this architecture are precipitated by SCI, wherein specific neuronal nodes form *de novo* maladaptive connections that permit and exacerbate the emergence of autonomic dysreflexia. This architecture has its foundation in Calca^ON^ neurons located in the dorsal root ganglia, which establish maladaptive projections to SC^Hoxa10::Zfhx3::Vsx2^ neurons in the lumbosacral spinal cord. In turn, an SCI provokes these neurons to propel axons to the *hemodynamic hotspot* located within the lower thoracic spinal cord, where they form *de novo* synaptic connections with SC^Hoxa7::Nfib::Vsx2^ neurons. These locally-projecting neurons establish connections onto Chat^ON^ sympathetic preganglionic neurons, which permit massive blood pressure elevations through the recruitment of neurons in splanchnic ganglia and subsequent alpha_1_ receptor activation throughout the dense splanchnic vasculature. The consequence of this aberrant neuronal architecture is the emergence of life-threatening autonomic dysreflexia.

## Competitive neuronal architectures converge on SC^Hoxa7::Nfib::Vsx2^ neurons

Our results demonstrate that SCI precipitates the formation of an aberrant neuronal architecture that exploits the natural connections from SC^Hoxa7::Nfib::Vsx2^ neurons onto sympathetic preganglionic neurons located in the *hemodynamic hotspot* to trigger autonomic dysreflexia. We reasoned that competitive engagement of the same neuronal subpopulations with an intervention that mediates beneficial as opposed to maladaptive reorganization of synaptic projections onto SC^Hoxa7::Nfib::Vsx2^ neurons could prevent the emergence of autonomic dysreflexia. Using molecular cartography, we recently demonstrated that epidural electrical stimulation (EES) of the lumbar spinal cord restores walking through the activation of locally-projecting Vsx2^ON^ neurons in the lumbar spinal cord^27,39^. We surmised that the same principle must exist in the lower thoracic spinal cord, since we previously showed that EES applied over this *hemodynamic hotspot* triggers robust pressor responses after SCI^19^. We thus hypothesized that EES applied over the lower thoracic spinal cord could compete with SC^Hoxa10::Zfhx3::Vsx2^ neurons to modulate SC^Hoxa7::Nfib::Vsx2^ neurons.

To identify the neuronal subpopulations engaged by EES applied to the lower thoracic spinal cord, we performed an additional comparative snRNA-seq experiment (**Supplementary Fig. 6a-c**). Concretely, we profiled neuronal nuclei from mice with SCI that had received EES over the lower thoracic spinal cord during 30 minutes. As we previously described in rats, non-human primates, and humans with SCI^19^, all the mice exhibited robust pressor responses when delivering EES over the lower thoracic spinal cord, referred to as the *hemodynamic hotspot* (**Fig. 3a-b**). High-quality transcriptional profiles were obtained from 21’098 nuclei (**Fig. 3c**). Integration of this dataset^66^ with our previous atlases^27,29,30,39,40^ and data from the experiments conducted on mice that were exposed to repetitive autonomic dysreflexia enabled us to identify and evaluate the same neuronal subpopulations. Cell type prioritization^39,41^ revealed that locally-projecting SC^Hoxa7::Nfib::Vsx2^ neurons exhibited the most pronounced transcriptional response across the compendium of neuronal subpopulations embedded in the lower thoracic spinal cord of mice that had received EES targeting the *hemodynamic hotspot* (**Fig. 3c, Supplementary Fig. 6d-e**).

**Fig. 3.**
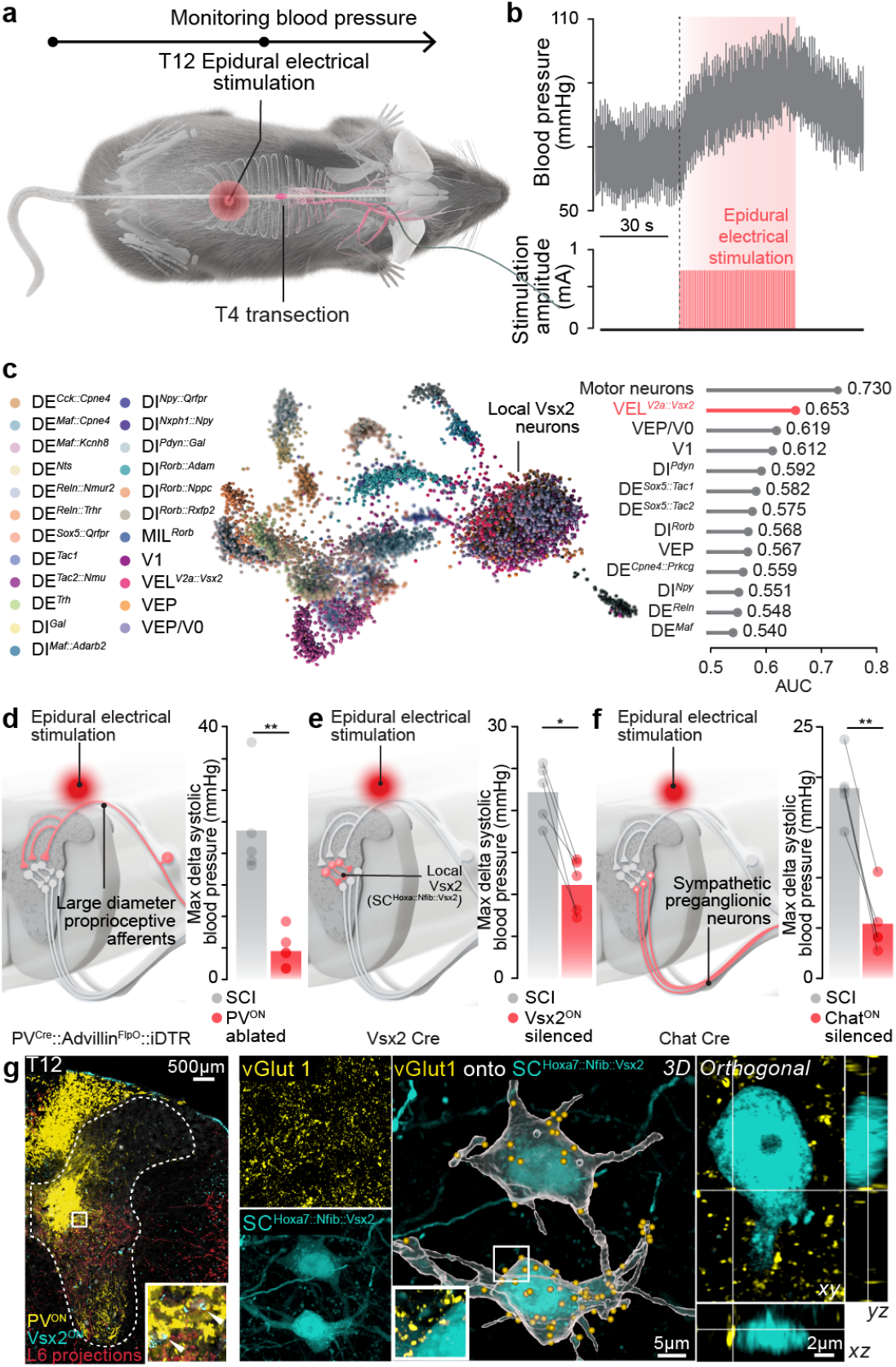
The neuronal architecture of epidural electrical stimulation (EES)-induced pressor responses. **a**, Schematic overview of experiments to trigger pressor responses with EES in mice with SCI. **b**, Pressor response induced by continuous (40 Hz) EES in a mouse with SCI. **c**, Uniform manifold approximation and projection (UMAP) visualization of 21,098 neuronal nuclei, colored by neuronal subpopulation identity. *Right*, Identification of perturbation-responsive neuronal subpopulations with Augur. **d-f**, Schematic overview of the successive nodes constituting the neuronal architecture though which EES applied over the low thoracic spinal cord induces pressor responses. **d**, EES-induced pressor responses before and after the ablation of PV^ON^ neurons located in the dorsal root ganglia in PV^Cre^::Advil^FlpO^::iDTR mice (n = 5; independent samples t-test; t = -5.41; p-value = 0.0043). **e**, EES-induced pressor responses before and after chemogenetic silencing of Vsx2^ON^ located in the lower thoracic spinal cord in Vsx2^Cre^ mice (n = 5; paired samples t-test; t = -4.21; p-value = 0.014). **f**, EES-induced pressor responses before and after chemogenetic silencing of Chat^ON^ neurons located in the lower thoracic spinal cord in Chat^Cre^ mice (n = 5; paired samples t-test; t = -7.07; p-value = 0.0021). **g**, Photomicrograph of the lower thoracic spinal cord demonstrating vGlut1^ON^ synaptic puncta and synaptic-like appositions from large-diameter afferent neurons onto SC^Hoxa7::Nfib::Vsx2^ neurons labelled with *in situ* hybridization (*Left*) or viral tract tracing (*Right*) in the lower thoracic spinal cord of PV^Cre^::Advil^FlpO^::tdTomato mice.

Since we previously found that Vsx2^ON^ neurons were recruited in response to EES in the lumbar spinal cord^27,39^, these comparative snRNA-seq experiments indicate that the principle through which EES recruits specific neuronal subpopulations is conserved across the thoracolum-bar spinal cord. Moreover, this observation nominates SC^Hoxa7::Nfib::Vsx2^ neurons as a convergence node that is not only recruited during autonomic dysreflexia but can also be engaged by the neuronal architecture that prevents orthostatic hypotension when the delivery of EES targets the *hemodynamic hotspot*.

The discovery of this intersection compelled us to expose the entire neuronal architecture activated by EES. Therefore, we conducted sequential anatomical and functional experi-ments that aimed to reconstruct the successive nodes involved in the pressor responses during the delivery of EES targeting the *hemodynamic hotspot*.

We previously showed that EES applied over the lower thoracic spinal cord activates afferent fibers in the posterior roots to trigger pressor responses^19^, and mounting evidence suggests that EES restores walking through the recruitment of large-diameter fibers where they bend to enter the spinal cord through the dorsal root entry zones^67,68^. We thus asked whether large-diameter afferent neurons act as the first node of the neuronal architecture that enables EES targeting the *hemodynamic hotspot* to trigger pressor responses (**Supplementary Fig 7a-b**).

To answer this question, we administered diphtheria toxin to PV^Cre^::Advil^FlpO^::iDTR mice and Calca^Cre^::Advil^FlpO^::iDTR mice, and applied EES targeting the *hemodynamic hotspot*. The ablation of PV^ON^ neurons in the dorsal root ganglia abolished pressor responses to EES, whereas the suppression of Calca^ON^ neurons failed to alter these pressor responses (**Fig. 3d, Supplementary Fig 7b-d**). We then verified that large-diameter afferents establish synaptic projections onto SC^Hoxa7::Nfib::Vsx2^ neurons in the lower thoracic spinal cord. To expose this connectome, we visualized large-diameter afferent fibers in PV^Cre^::Advil^FlpO^::Ai9^(RCL-tdT)^ mice and confirmed that these fibers established vGlut1^ON^ synaptic-appositions onto SC^Hoxa7::Nfib::Vsx2^ neurons (**Supplementary Fig. 7e-f**). The dense projections from PV^ON^ neurons onto SC^Hoxa7::Nfib::Vsx2^ neurons contrasted with the absence of projections from PV^ON^ neurons onto Chat^ON^ neurons in the intermediolateral column of the lower thoracic spinal cord, both before or after SCI (**Supplementary Fig. 7g-h**).

Since SC^Hoxa7::Nfib::Vsx2^ neurons underwent the greatest transcriptional perturbation following EES targeting the *hemodynamic hotspot* and receive direct projections from large-diameter afferent fibers that are recruited by EES, we anticipated that SC^Hoxa7::Nfib::Vsx2^ neurons act as the second node in the neuronal architecture that enables EES to trigger pressor responses. To expose the necessary role of SC^Hoxa7::Nfib::Vsx2^ neurons, we infused AAV5-hSyn-DIO-hM4Di-mCherry into the lower thoracic spinal cord of Vsx2^Cre^ mice. Inactivation of SC^Hoxa7::Nfib::Vsx2^ neurons blunted the pressor responses triggered by EES (**Fig. 3e, Supplementary Fig. 7i-j**).

We surmised that Chat^ON^ sympathetic preganglionic neurons act as the third and final node in the neuronal architecture that enables EES targeting the *hemodynamic hotspot* to trigger pressor responses. Indeed, chemogenetic inactivation of Chat^ON^ neurons located in the lower thoracic spinal cord abolished pressor responses triggered by EES (**Fig. 3f, Supplementary Fig. 7k-l**).

Together these results uncovered the neuronal architecture that enables EES targeting the *hemodynamic hotspot* of the lower thoracic spinal cord to trigger pressor responses. EES directly recruits large diameter afferents to activate SC^Hoxa7::Nfib::Vsx2^ neurons, which project to Chat^ON^ sympathetic preganglionic neurons to elicit pressor responses through the activation of ganglionic neurons, and subsequent alpha_1_ receptor activation to induce vasoconstriction.

Finally, we surmised that for the neuronal architecture enabling EES to elicit pressor responses to compete with the neuronal architecture that triggers autonomic dysreflexia, these two architectures must intersect on the same SC^Hoxa7::Nfib::Vsx2^ neurons in the lower thoracic spinal cord.

To expose this anatomical and functional intersection, we first infused AAV5-hSyn-eGFP (enhanced green fluorescent protein) into the lumbosacral spinal cord of PV^Cre^::Advil^FlpO^::Ai9^(RCL-tdT)^ mice with SCI coupled to labeling of SC^Hoxa7::Nfib::Vsx2^ neurons. We found that the same SC^Hoxa7::Nfib::Vsx2^ neurons received direct projections from large-diameter afferents emanating from PV^ON^ neurons located in dorsal root ganglia, and from axons projecting from the lumbosacral spinal cord after SCI (**Fig. 3g**). Second, we aimed to confirm that these two distinct axonal projections were able to regulate the activity of the same SC^Hoxa7::Nfib::Vsx2^ neurons. We conducted single-unit recordings of optogenetically-identified SC^Hoxa7::Nfib::Vsx2^ neurons in the lower thoracic spinal cord in response to autonomic dysreflexia and EES. We found that both paradigms elicited short-latency responses in the recorded SC^Hoxa7::Nfib::Vsx2^ neurons that were compatible with a direct activation of the same SC^Hoxa7::Nfib::Vsx2^ neurons.

## Neuronal architecture competition to reverse autonomic dysreflexia

The intersection between the neuronal architecture that enables EES targeting the *hemodynamic hotspot* to elicit pressor responses and the neuronal architecture that triggers autonomic dysreflexia opened the intriguing possibility that the sustained modulation of SC^Hoxa7::Nfib::Vsx2^ neurons with EES could compete with the detrimental activity emanating from aberrant axonal projections of SC^Hoxa10::Zfhx3::Vsx2^ neurons, ultimately reversing autonomic dysreflexia.

To test this possibility, we subjected mice to autonomic neurorehabilitation, which consisted of daily sessions during which EES targeting the *hemodynamic hotspot* was delivered over the lower thoracic spinal cord during the course of one month (**Fig. 4a, Supplementary Fig. 8a**). Autonomic neurorehabilitation abolished autonomic dysreflexia in all tested mice (**Fig. 4b, Supplementary Fig. 8b**).

**Fig. 4.**
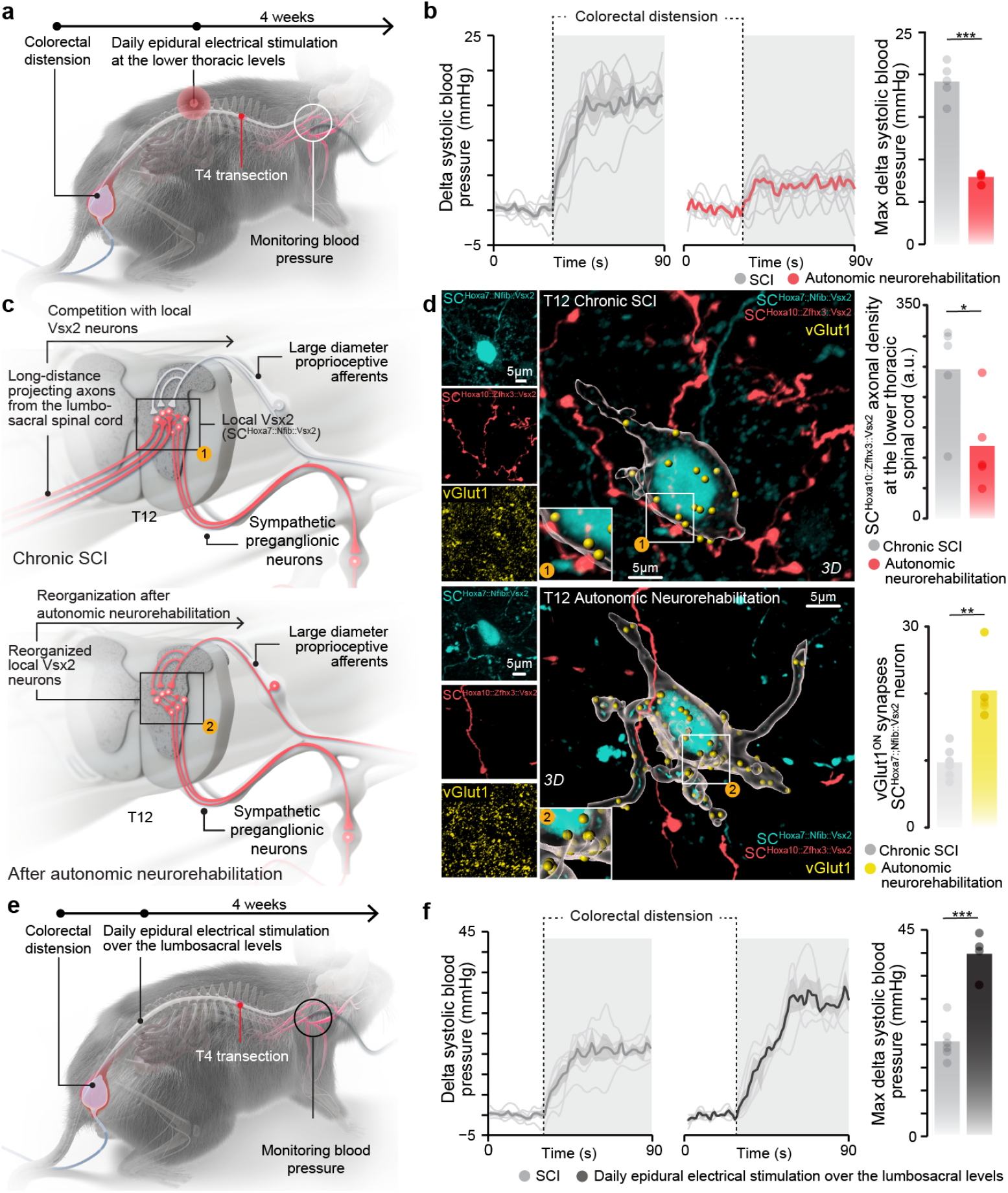
Competitive neuronal architectures converge on SC^Hoxa7::Nfib::Vsx2^ neurons. **a**, Schematic overview of autonomic neurorehabilitation and paradigm to quantify the severity of autonomic dysreflexia. **b**, Pressor responses (*Left* ; individual mice and mean trace) and severity of autonomic dysreflexia (*Right*) in 5 with chronic SCI and 5 mice that underwent autonomic neurorehabilitation for 4 weeks, starting 1 week after SCI (independent samples t-test; t = -7.45; p-value = 0.00056). **c**, Schematic overview illustrating the competitive (overlapping) neuronal architectures of autonomic dysreflexia and EES-induced pressor responses, and their rearrangement after autonomic neurorehabilitation. **d**, vGlut1^ON^ synaptic puncta and synaptic-like appositions from SC^Hoxa10::Zfhx3::Vsx2^ neurons onto SC^Hoxa7::Nfib::Vsx2^ neurons in in mice with SCI and mice with SCI that underwent autonomic neurorehabilitation. (*top*) Bar plots reporting the mean density of axonal projections from SC^Hoxa10::Zfhx3::Vsx2^ neurons in the thoracic spinal cord in mice with SCI and mice with SCI that underwent autonomic neurorehabilitation (n = 5; independent samples t-test; t = 2.51; p-value = 0.0369). (*bottom*) (bottom) Bar plots reporting the mean number of vGlut1^ON^ synaptic puncta apposing SC^Hoxa7::Nfib::Vsx2^ neurons (n = 5; independent samples t-test; t = 4.44; p-value = 0.0055). **e**, Schematic overview of experiments in which EES was applied daily over the lumbosacral spinal cord of mice with SCI, and paradigm to quantify the severity of autonomic dysreflexia. **f**, As in **b**, for mice with SCi that were subjected to the daily application of EES over the lumbosacral spinal cord (n = 5; independent samples t-test; t = 5.82; p-value = 0.00070).

We suspected that the suppression of autonomic dysreflexia would emerge from the competitive advantage of large-diameter fiber projections over aberrant axonal projections from SC^Hoxa10::Zfhx3::Vsx2^ neurons to form synaptic connections with SC^Hoxa7::Nfib::Vsx2^ neurons (**Fig. 4c**). To expose this adversarial anatomy, we labeled SC^Hoxa7::Nfib::Vsx2^ and SC^Hoxa10::Zfhx3::Vsx2^ neurons with intersectional viral tracing strategies. We then quantified the relative density of vGlut1^ON^ synapses emanating from PV^ON^ large-diameter afferent fibers compared to the density of axonal projections from SC^Hoxa10::Zfhx3::Vsx2^ neurons. Autonomic rehabilitation reduced the number of aberrant projections from SC^Hoxa10::Zfhx3::Vsx2^ neurons onto SC^Hoxa7::Nfib::Vsx2^ neurons, while concomitantly augmenting the density of vGlut1^ON^ synaptic appositions onto SC^Hoxa7::Nfib::Vsx2^ neurons embedded in the *hemodynamic hotspot* (**Fig. 4d, Supplementary Fig. 8c-d**).

## Mistargeted stimulation exacerbates autonomic dysreflexia

Recent case studies reported transient increases in blood pressure in response to EES applied over the lumbosacral spinal cord of people with SCI^69–74^, and that the long-term delivery of EES over this location led to improvements in resting hypotension and orthostatic hypotension in these individuals. Since a paucity of neurons involved in the control of blood pressure reside in this region of the spinal cord, the activation and reinforcement of the neuronal architecture that causes autonomic dysreflexia is the most likely mechanism to account for these observations. Therefore, we hypothesized that the daily activation of SC^Hoxa10::Zfhx3::Vsx2^ neurons in response to EES applied over the lumbosacral spinal would provide a competitive advantage to vGlut2^ON^ synapses emanating from these neurons and projecting onto SC^Hoxa10::Zfhx3::Vsx2^ neurons compared to vGlut1^ON^ synapses received from local PV^ON^ large-diameter afferent fibers, and that ultimately, this shift in the synaptic innervations of SC^Hoxa10::Zfhx3::Vsx2^ neurons would exacerbate autonomic dysreflexia.

To test this hypothesis, we subjected a new cohort of mice to daily exposure to EES, but instead of delivering EES over the lower thoracic spinal cord to target the *hemodynamic hotspot*, we applied EES over the lumbosacral spinal cord for 30 minutes during the course of one month (**Fig. 4e, Supplementary Fig. 8e**). This mistargeted stimulation doubled the severity of autonomic dysreflexia in all tested mice (**Fig. 4f, Supplementary Fig. 8f**). As anticipated, we observed a concomitant increase in the density of axonal projections from SC^Hoxa10::Zfhx3::Vsx2^ neurons onto SC^Hoxa7::Nfib::Vsx2^ neurons, combined with a decrease in the density of vGlut1^ON^ synapses from PV^ON^ large-diameter afferent fibers (**Supplementary Fig. 8g-h**).

The dramatic exacerbation of autonomic dysreflexia triggered by the mistargeted delivery of EES demonstrates that attempts to restore hemodynamic stability with EES applied over the lumbosacral spinal cord are not only less efficacious than EES targeting the *hemodynamic hotspot* located in the lower thoracic spinal cord, but also proved unsafe, since the mechanism by which hemodynamics can be modulated when stimulating this region is by triggering and reinforcing the severity of life-threatening autonomic dysreflexia.

## Longitudinal monitoring of autonomic neurorehabilitation

While autonomic neurorehabilitation suppressed autonomic dysreflexia in mice, we recognized that the longitudinal monitoring of the safety and efficacy of this treatment over a long period of time would be important to inform the design of a therapy for people living with SCI.

We therefore leveraged our chronic rat model of high-thoracic contusion SCI that enables 24/7 monitoring of hemodynamic parameters without the constraints of tethered electronics^19,75^(**Fig. 5a, Supplementary Fig. 9a**). We surgically inserted an electronic dura mater (*e-dura*) implant with an optimized configuration of electrodes to target the dorsal root entry zones projecting to the last three thoracic segments of the spinal cord, which coincide with the *hemodynamic hotspot*. Closed-loop adjustment of EES amplitudes with a proportional-integral controller enabled the maintenance of blood pressure to a predefined target^19^ (**Fig. 5b**). We previ-ously established the efficacy of this technology for the treatment of orthostatic hypotension, which we named the *neuroprosthetic baroreflex*^19^.

**Fig. 5.**
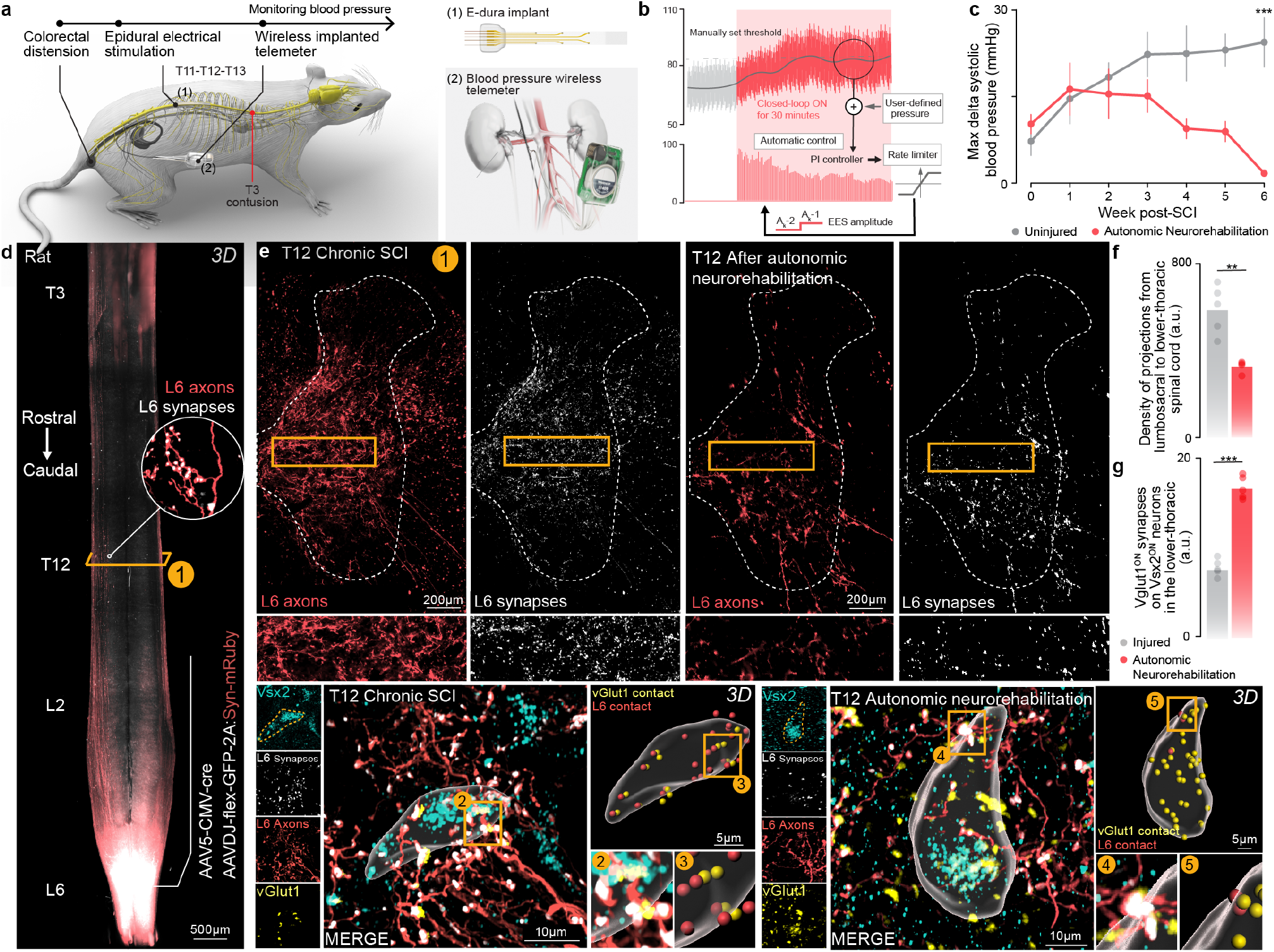
Longitudinal monitoring of autonomic neurorehabilitation in rats. **a**, Schematic overview of autonomic neurorehabilitation in rats. A wireless telemetry system was implanted chronically to acquire longitudinal recordings of haemodynamic parameters. An electronic dura mater (*e-dura*) designed to target the dorsal roots projecting to the T11, T12, and T13 spinal segments was then implanted over the *hemodynamic hotspot* to regulate blood pressure. **b**, Augmentation of systolic blood pressure within a target range using a proportional integral (PI) controller that adjusts the amplitude of EES in closed-loop. **c**, Line graph reporting the severity of autonomic dysreflexia that was assessed weekly using colorectal distension in rats with SCI and rats with SCI that were undergoing autonomic neurorehabilitation. Raw data and statistics provided in **Supplementary Table 1**. **d**, Whole spinal cord visualization of projections from neurons located in the lumbosacral spinal cord that establish neurons into the lower thoracic spinal cord. **e**, Axonal projections and synaptic puncta from neurons located in the lumbosacral spinal cord and projecting to the lower thoracic spinal cord, shown in rats with SCI and rats with SCI that underwent autonomic neurorehabilitation. **f**, Density of axonal projections from lumbosacral neurons in the thoracic spinal cord before and after autonomic neurorehabilitation (n = 5; independent samples t-test; t = -4.03; p-value = 0.0081). **g**, Synaptic-like appositions from neurons located in the lumbosacral spinal cord onto SC^Hoxa7::Nfib::Vsx2^ neurons combined with the labelling of vGlut1^ON^ synaptic puncta form large-diameter afferent fibers in a rat with chronic SCI and a rat with SCI that underwent autonomic neurorehabilitation. **h**, Bar plots reporting the mean density of vGlut1^ON^ synaptic puncta onto SC^Hoxa7::Nfib::Vsx2^ neurons in rats with SCI and rats with SCI that underwent autonomic neurorehabilitation (n = 5; independent samples t-test; t = 12.71; p-value = 2.78e-06).

We exposed rats with SCI to daily sessions of autonomic neurorehabilitation. During these sessions, we configured the closed-loop controller to augment blood pressure to a predefined target. Careful monitoring of hemodynamics during autonomic neurorehabilitation revealed that blood pressure was maintained within and did not exceed the defined target range, indicating that EES targeting the *hemodynamic hotspot* did not trigger autonomic dysreflexia (**Fig. 5b**). Moreover, blood pressure decreased as soon as EES was switched off.

We conducted formal weekly hemodynamic assessments to quantify the severity of autonomic dysreflexia (**Fig. 5c, Supplementary Fig. 9c**). These assessments revealed that autonomic dysreflexia vanished in all tested rats. Rats with chronic SCI exhibited an increase in the number of axonal projections emanating from neurons located in lumbosacral segments that formed synaptic appositions with Vsx2^ON^ neurons in the lower thoracic spinal cord, comparable to observations in mice (**Fig. 5d, Supplementary Fig. 9d-g**). Autonomic neurorehabilitation reversed this aberrant connectome, and in turn augmented the density of vGlut1^ON^ synapticappositions from PV^ON^ neurons onto Vsx2^ON^ neurons (**Fig. 5d, Supplementary Fig. 9h-k**).

These results reinforced the hypothesis that autonomic neurorehabilitation is safe and enables large-diameter afferents to compete for the innervation of Vsx2^ON^ neurons located in the *hemodynamic hotspot*, and ultimately, to overcome the aberrant neuronal architecture responsible for the emergence of autonomic dysreflexia. The consequence is the reversal of autonomic dysreflexia.

## Clinical necessity of therapies for autonomic neurorehabilitation

While mice and rat experiments documented the safety and efficacy of EES targeting the *hemodynamic hotspot* to reduce the severity of autonomic dysreflexia, the neurosurgical intervention necessary to deploy this therapy must be weighed against the risks and benefits of the procedure. Establishing this balance requires an understanding of the prevalence, symptomatology, and effectiveness of current management strategies.

Consequently, we leveraged the Spinal Cord Injury Community Survey (SCICS), which includes self-reported information on the presence of symptoms of autonomic dysreflexia, and demographic information about the level of SCI (n = 1,479)^76,77^. These data revealed that 82% of individuals with tetraplegia had been told by a medical practitioner that they present autonomic dysreflexia. We found that only 30% of the individuals were being treated for autonomic dysreflexia, yet 98% of these treated individuals still experienced symptoms (**Fig. 6a, Supplementary Fig. 10a**).

**Fig. 6.**
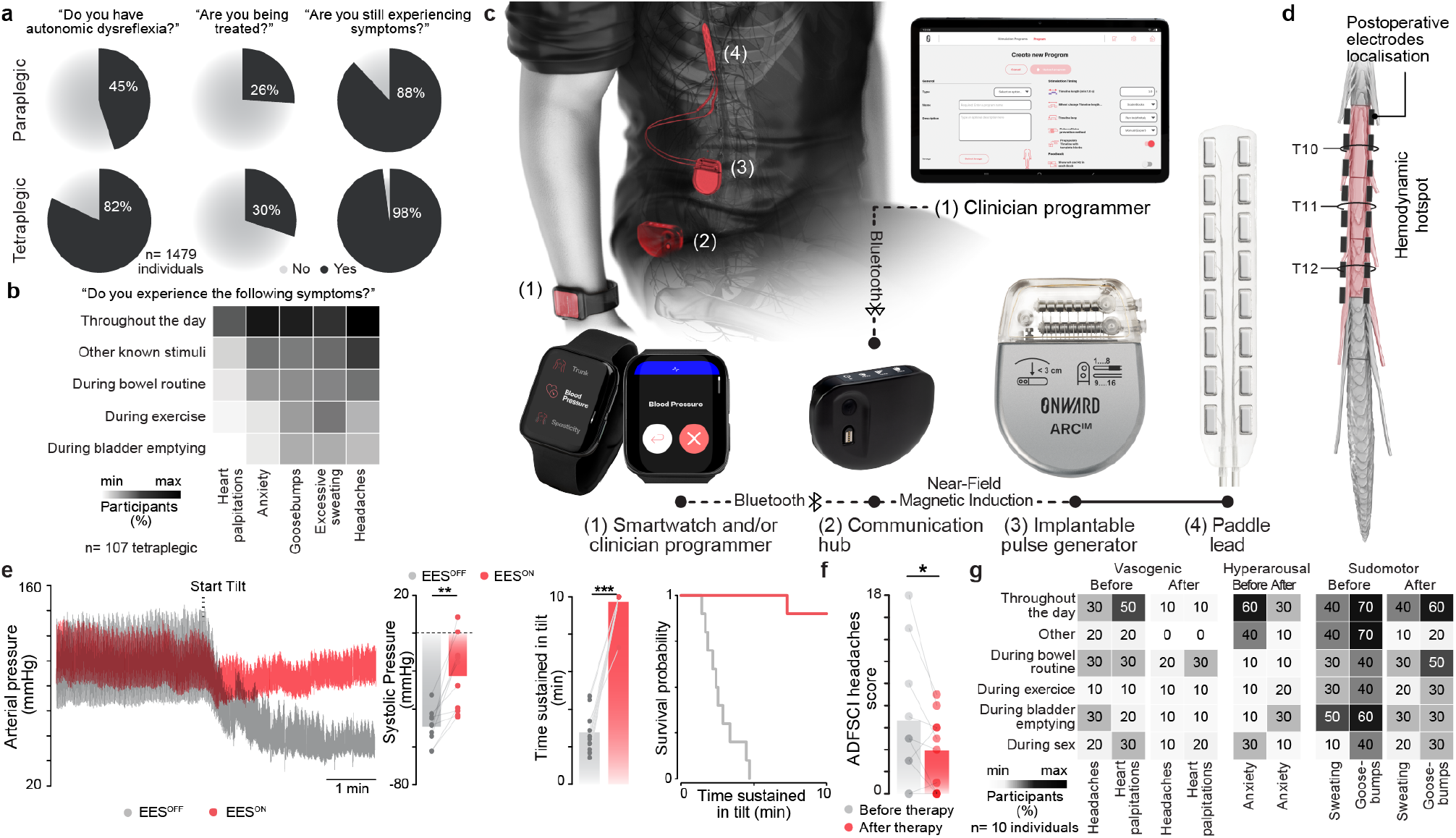
Reduced severity of autonomic dysreflexia in people with chronic SCI following autonomic neurorehabilitation. **a**, Prevalence of autonomic dysreflexia and management efficacy quantified in 1,479 individuals with SCI from the Rick Hansen Spinal Cord Injury Registry^76,77^. **b**, Percentage of individuals with tetraplegia experiencing each symptom of autonomic dysreflexia scored in the Autonomic Dysfunction following SCI (ADFSCI) across various daily activities (n = 107). **c**, Implantable system to regulate blood pressure with EES, including a paddle lead with optimized electrode configurations to target the dorsal roots projecting to the hemodynamic hotspot, an implantable pulse generator, communication hub, and external smartwatch to operate the various programs of the therapy. **d**, Post-operative reconstruction of the final position of the electrodes following the implantation of the paddle lead. **e**, Changes in blood pressure from a representative participant during an orthostatic challenge without EES and with continuous EES applied over the *hemodynamic hotspot*. The bar plots report the average drop in systolic blood pressure during orthostatic challenge (n = 10, paired samples two-tailed t-test; t = 4.7774 ; p = 0.00101) and average tilt duration without EES and with EES applied over the *hemodynamic hotspot* (n = 10, paired samples two-tailed t-test; t = 14.33100 ; p < 0.001). Kaplan-Meier plot of exposure status to time, segregated by the presence or absence of EES. **f**, ADFSCI headaches score before implantation and after at least 6 months but up to 2 years after implantation of the system and daily use to regulate blood pressure (n = 10, paired samples one-tailed t-test; t = 2.21 ; p =0.027). **g**, Percentage of individuals (n = 10) experiencing each symptom described in the ADFSCI autonomic dysreflexia section before implantation (top) and after 6 months of ARC^IM^ Therapy (bottom).

To further characterize the pattern of hypertensive-related symptoms in individuals with SCI, we next surveyed a diverse sample of 254 individuals with SCI from 50 different countries, and included responses to the Autonomic Dysfunction Following Spinal Cord Injury (ADFSCI) rating scale. When considering individuals with tetraplegia, we found that 79% of these individuals experienced symptoms of autonomic dysreflexia (**Fig. 6b, Supplementary Fig. 10b**). These symptoms were primarily characterized by vasogenic manifestations including headaches and heart palpitations, sudomotor manifestations including goose bump and sweat-ing, as well as anxiety (**Fig. 1c**).

This symptomatology combined with the risk of life-threatening episodes of autonomic dysreflexia and the absence of adequate therapeutic management justified evaluating the impact of autonomic neurorehabilitation on the severity of autonomic dysreflexia in humans living with chronic SCI.

## Clinical validation of autonomic neurorehabilitation

We next aimed to conduct a preliminary clinical evaluation to assess whether the long-term application of EES targeting the *hemodynamic hotspot* to reduce hypotensive symptomatology also reduces the severity of autonomic dysreflexia in humans with chronic SCI. To carry out this assessment, we leveraged four ongoing clinical studies (**STIMO-HEMO** [NCT04994886, CHUV, Lausanne, Switzerland], **HEMO** [NCT05044923, University of Calgary, Calgary, Canada], **HemON** [NCT05111093, CHUV, Lausanne, Switzerland] and **HemON-NL** 05941819, Sint Maartenskliniek, Ubbergen, Netherlands]) focused on the development of purpose-built technologies to restore hemodynamic stability based on EES targeting the *hemodynamic hotspot*.

These studies enrolled patients with cervical SCI who presented with medically-refractory orthostatic hypotension. When exposed to orthostatic challenges during a tilt-table test, all the participants exhibited a rapid decline in blood pressure that required the early termination of the tilt-test. Following this verification of their eligibility, the participants underwent a neurosurgical intervention to implant an electrode array targeting the dorsal root entry zones innervating the last three thoracic segments and a neurostimulator to deliver EES. Post-operative evaluations confirmed that EES targeting the *hemodynamic hotspot* elicited robust pressor responses that reduced the severity of orthostatic hypotension. The participants then learned how to operate the therapy in order to regulate their blood pressure throughout the day for up to two years.

While these studies focused on the long-term reduction of hypotensive symptomatology, the severity of autonomic symptoms was assessed concurrently with the ADFSCI rating scale^78^. Consequently, the context of these studies allowed us to assess how the symptomatology related to autonomic dysreflexia evolved in nine participants.

Quantification of the autonomic dysreflexia subscore within the ADFSCI revealed a decrease in symptomatology related to autonomic dysreflexia after long-term use of EES targeting the *hemodynamic hotspot*. Secondary analyses of vasogenic and sudomotor manifestations revealed that the long-term use of EES targeting the *hemodynamic hotspot* led to a reduction in headaches and heart palpitations, which are the main vasogenic symptoms of autonomic dysreflexia. In stark contrast, sudomotor manifestations and anxiety remained unchanged from baseline.

These clinical outcomes provide preliminary evidence that the daily application of EES targeting the *hemodynamic hotspot* reduces the severity of autonomic dysreflexia in humans with chronic SCI, as quantified functionally and anatomically in mice and rats.

## Discussion

Here, we exposed the complete *de novo* neuronal architecture that develops after SCI to cause autonomic dysreflexia. This architecture incorporates locally-projecting, Vsx2-expressing neurons located in the lower thoracic spinal cord that are also embedded in the neuronal architecture that enables EES targeting the *hemodynamic hotspot* to achieve safe increases of blood pressure. Since these adversarial architectures converged onto a single neuronal subpopulation, their relative activation determined the prevailing architecture in this competitive interaction.

Exposing these adversarial architectures allowed us to design a safe intervention that reversed autonomic dysreflexia in mice, rats, and humans with SCI by applying EES targeting the *hemodynamic hotspot* located in the lower thoracic spinal cord. Conversely, the mistargeted application of EES over the lumbosacral spinal cord reinforced the anatomical and functional connectivity of the neuronal architecture responsible for autonomic dysreflexia, which augmented the severity of these symptoms. These mechanisms establish that the long-term delivery of EES over the lumbosacral spinal cord to stabilize hemodynamic stability is contraindicated for people who suffer from autonomic dysreflexia after SCI^69–74^.

The involvement of Vsx2-expressing neurons in the emergence of autonomic dysreflexia exposed the unique properties of these neurons in the regulation of neurological functions and circuit reorganization processes after neurological disorders and neurotraumas. Indeed, we recently demonstrated that Vsx2-expressing neurons located in the spinal cord are inherently primed to overexpress circuit reorganization gene programs—thus becoming *recovery-organizing* neurons after SCI that guide the spontaneous and therapeutically-enhanced recovery of walking after both incomplete and complete SCI^27,29,30^. Here, we extend the role of these cells by demonstrating that since Vsx2-expressing neurons are primed to reorganize in response to injury, but are lacking molecular cues to direct their reorganization after severe SCI^79^, they can be naturally coerced towards undirected and aberrant neuronal architectures that drive the emergence of maladaptive neurological functions.

Autonomic dysreflexia is a life-threatening medical condition that can lead to stroke, heart attack, and death^1–7^. Consequently, people living with SCI and medical practitioners are taught to identify warning signs such as headaches, sweating, and goosebumps, since these signs inform on the presence of a noxious stimulus that is triggering autonomic dysreflexia, and must therefore be localized to resolve the ongoing hypertensive episode as quickly as possible^80^. Analysis of self-reported symptoms in patients from four clinical studies showed that EES targeting the *hemodynamic hotspot* reduced headaches and heart palpitations, which are both directly related to dangerous increases in blood pressure. Instead, this therapy had no detectable impact on the other warning signs such as goosebumps and sweating that could still inform on the presence of noxious stimulus. The most likely explanation for this dichotomy is the peak concentration of sympathetic preganglionic neurons responsible for hemodynamic regulation within the antepenultimate segment of the thoracic spinal cord^19,81^, which is directly targeted by EES. Instead, the sympathetic preganglionic neurons triggering symptoms such as sweating and goosebumps are distributed uniformly throughout the thoracic spinal cord and are consequently poorly targeted by EES restricted to the lower thoracic spinal cord.

We took advantage of ongoing clinical trials to expose the potential for EES targeting the *hemodynamic hotspot* to reduce autonomic dysreflexia in humans with SCI. However, these trials were not designed to demonstrate the safety and efficacy of this therapy, and consequently, the next steps must include the assessment of the safety and efficacy of EES targeting the *hemodynamic hotspot* to reduce autonomic dysreflexia in a pivotal clinical trial.

## Materials and Methods

### Mouse and rat models

Adult male or female C57BL/6 mice (15-25 g body weight, 8-15 weeks of age) or transgenic mice were used for all experiments. Vglut2^Cre^ (Jackson Laboratory 016963), Vsx2^Cre^ (MMMRRC 36672, also called Chx10^Cre^), Chat^Cre^, Vgat^Cre^, Ai65^(RCFL-tdT)^ (Jackson Laboratory 021875), Parvalbumin (PV)^Cre^ (Jackson Laboratory 017320), Advillin^FlpO^ (a gift from V. Abraira), iDTR, Calca^Cre^ transgenic mouse strains were bred and maintained on a mixed genetic background (C57BL/6). Adult female Lewis rats (180–220 g body weight, 14–30 weeks of age) were used for the rat experiments. Housing, surgery, behavioral experiments and euthanasia were all performed in compliance with the Swiss Veterinary Law guidelines. Manual bladder voiding and all other animal care was performed twice daily throughout the entire experiment. All procedures and surgeries were approved by the Veterinary Office of the Canton of Geneva (Switzerland; authorization GE67).

### Viral vectors and vector production

Viruses used in this study were either acquired commercially or produced at the EPFL core facility. The following AAV plasmids were used and detailed sequence information is available as detailed or upon request: AAVDJ-hSyn-flex-mGFP-2A-synaptophysin-mRuby^33^ (Stanford Vector Core Facility, reference AAV DJ GVVC-AAV-100), AAV9-CMV-Cre (Vector Biolabs 7014), AAV5-hSyn-eGFP (Addgene 50465-AAV5), AAV5-Syn-flex-ChrimsonR-tdT (Addgene 62723), AAV5-hSyn-DIO-hm4D^31^ (Gi)-mCherry (Addgene 44362), AAV5-CAG-flex-tdTomato (a gift from S. Arber), AAV5-hSyn-Con/Fon-eYFP (Addgene 55650) and rAAV2-EF1a-DIO-Flpo (Addgene 87306).

### Spinal cord injury (SCI) models

For mouse spinal cord injuries, a laminectomy was performed on the T4 vertebra to expose the T4 spinal segment. Complete transections were performed using angled microscissors. Rat spinal cord injuries were performed according to our previously published work^19^. In brief, a laminectomy was performed on the T3 vertebra to expose the T3 spinal segment. Following this the rat was transferred to the Infinite-Horizons (IH-0400 Impactor, Precision Systems and Instrumentation LLC) impactor^19^ stage, where the T2 and T4 spinous processes were securely clamped using modified Allis forceps^19^. The rat was stabilized on the platform and the impactor tip (2.5 mm) was properly aligned using a three-dimensional coordinate system moving platform. The IH system was set to deliver an impact force of 400 kdyn, with a 5 s dwell time^19^. Analgesia (buprenorphine, Essex Chemie AG, 0.01–0.05 mg per kg, subcutaneously) and antibiotics (amoxicillin 200 mg per 4 ml, Sandoz, 200 mg l^1^ ad libitum) were provided for 3 and 5 days after surgery, respectively. Bladders were manually expressed twice a day until the end of the experiment^82^.

### Autonomic neurorehabilitation

In mice, we delivered EES with conventional stimulation protocols consisted of continuous EES delivered at 50 Hz with 5 ms pulses at 100–150 *µ*A (2100 Isolated Pulse Stimulator, A-M Systems). Mice underwent autonomic neurorehabilitation consisting of EES applied for 30 minutes each day for 4 weeks, starting 1 week after SCI. In rats, we applied a closed-loop controlled stimulation using a proportional integral (PI) controller that adjusts the amplitude of traveling EES waves over the three *hemodynamic hotspot*. We applied a +10 mmHg systolic blood pressure target from the baseline acquired at the beginning of each autonomic neurorehabilitation session^19^. Rats received 30 minutes of closed-loop controlled stimulation for 6 weeks, starting one week after SCI.

### Neuron-specific ablation, chemogenetics and optogenetics

For ablation experiments with the diphtheria toxin, we used PV^Cre^::Advilin^FLPo^::iDTR and Calca^Cre^::Advillin^FLPo^::iDTR mice. Four weeks after the spinal cord injuries (T4 spinal level complete transection), mice received intraperitoneal injections of diphtheria toxin (Sigma, D0564) diluted in saline (100 *µ*gkg^1^) to target respectively PV^ON^ or Calca^ON^ neurons. Mice were tested two weeks post-injection. To manipulate the activity of Vglut2^ON^, Vgat^ON^ neurons, AAV5-hSyn-DIO-hm4D was infused in either the lower thoracic spinal cord (T11-T13) or lumbosacral spinal cord (L5-S1) of either Vglut2^Cre^ or Vgat^Cre^ mice prior to performing the spinal cord injury. To manipulate Chat^ON^ neuronal activity, AAV5-hSyn-DIO-hm4D was infused in the lower thoracic spinal cord (T11-T13) of Chat^Cre^ mice prior to performing the spinal cord injury. To ma-nipulate ^SCHoxa7::Nfib::Vsx2^ or SC ^Hoxa10::Zfhx3::Vsx2^neurons, AAV5-hSyn-DIO-hm4D was infused in either the lower thoracic spinal cord (T11-T13) or lumbosacral spinal cord (L5-S1), respectively, in Vsx2^Cre^ mice prior to performing the spinal cord injury. After four weeks, autonomic dysreflexia or EES-induced pressor response were assessed before and between 30–45 min after intraperitoneal injections of 5 mgkg^1^ clozapine N-oxide (CNO) (Carbosynth, CAS: 34233-69-7, suspended in 2% DMSO in saline). To optogenetically manipulate Vsx2^ON^ neurons, AAV5-Syn-flex-Chrimson was infused in either the lower thoracic spinal cord (T11-T13) and the lumbosacral spinal cord (L5-S1), in Vsx2^Cre^ mice prior to performing the spinal cord injury. After six weeks, laminectomies were made over T11/T12/T13 and L5/L6/S1 spinal segments. 5-ms pulses were delivered at 50 Hz from a 635 nm laser (LaserGlow Technologies LRD-0635-PFR-00100-03). Laser light was delivered to the surface of the spinal cord through a fibre optic cable attached to 400 *µ*m, 0.39 NA cannula with a 5 mm tip (Thorlabs). Optical power was set to 2.35 mW at the tip.

### Spinal injections for exclusive labeling of Vsx2^ON^ neurons

To exclusively label Vsx2^ON^ neurons in the lumbosacral spinal cord with long-distance projections to the lower-thoracic region (SC^Hoxa10::Zfhx3::Vsx2^), we leveraged Boolean logic viral strategies^65^. Partial laminectomies were made over the T11/T12/T13 and L5/L6/S1 spinal segments of Vsx2^Cre^ mice. Two sets of bilateral injections of AAV5-hSyn-Con/Fon-eYFP (Addgene 55650)^65^ were made over the L5/L6/S1 spinal segments (0.25 *µ*l per injection) at a depth of 0.6 mm below the dorsal surface and separated by 1 mm. Two sets of bilateral injections of rAAV2-EF1a-DIO-Flpo (Addgene 87306) were made over the T11/T12/T13 spinal segments (0.15 *µ*l per injection) at two depths (0.8 mm and 0.4 mm below the dorsal surface) and separated by 1 mm. Animals were perfused four weeks later. To label lower-thoracic Vsx2^ON^ neurons (SC^Hoxa7::Nfib::Vsx2^), two sets of bilateral injections of AAV-flex-tdTomato were made over T11/T12/T13 spinal levels (0.15 *µ*l per injection) at two depths (0.8 mm and 0.4 mm below the dorsal surface) and separated by 1 mm.

### Perfusions

Animals were perfused at the end of the experiments. Animals were deeply anesthetized by an intraperitoneal injection of 0.2 m sodium pentobarbital (50 mg/ml). Animals were transcardially perfused with Phosphate buffered saline (PBS) followed by 4% paraformaldehyde in PBS. Tissues were removed and post-fixed overnight in 4% paraformaldehyde before being transferred to PBS or cryoprotected in 30% sucrose in PBS.

### Immunohistochemistry

Immunohistochemistry was performed as described previously^83–85^. Perfused post-mortem tissue was cryoprotected in 30% sucrose in PBS for 48 hours before being embedded in cryomatrix (Tissue Tek O.C.T, Sakura Finetek Europe B.V.) and freezing. 30 *µ*m thick transverse or horizontal sections of the spinal cord were cut on a cryostat (Leica), immediately mounted on glass slides and dried or in free floating wells containing PBS + 0.03% sodium azide. The sections were incubated with following primary antibody diluted in blocking solution at room temperature overnight: rabbit anti-cFos (1:500), chicken anti-vGlut1 (1:500), goat anti-Chat (1:100), rabbit anti-Chx10 (now known as Vsx2) (1:500, Synaptic Systems Gmbh). Fluorescent secondary antibodies were conjugated to Alexa 488 (green), or Alexa 405 (blue), or Alexa 555 (red), or Alexa 647 (far red) (ThermoFisher Scientific, USA). Nuclear stain: 4’,6’-diamidino-2-phenylindole dihydrochloride (DAPI; 2ng/ml; Molecular Probes). Sections were imaged digitally using a slide scanner (Olympus VS-120 Slide scanner) or confocal microscope (Zeiss LSM880 + Airy fast module with ZEN 2 Black software (Zeiss, Oberkochen, Germany). Images were digitally processed using ImageJ (ImageJ NIH) software or Imaris (Bitplane, v.9.8.2).

### Fluorescence ^in^ situ hybridization (FISH)

We performed *in situ* hybridization of cell type markers and using RNAscope (Advanced Cell Diagnostics). Lists of putative marker genes were obtained from snRNA-seq data, as described below, and crossreferenced against a list of validated probes designed and provided by Advanced Cell Diagnostics, Inc. Probes were obtained for the following genes: Chat, catalog no. 408731; and Vsx2, catalog no. 438341, Slc17a6, catalog no. 319171; Slc6a5, catalog no. 409741.We then generated 12 *µ*m cryosections from fixed-frozen spinal cords as previously described 91 and performed FISH for each probe according to the manufacturer’s instructions, using the RNAscope HiPlex kit (cat no. 324106).

### iDISCO+

Mice underwent a 90-min colorectal distension protocol (30 s inflate then 60 s deflate repeatedly)^3^ and were perfused^22,23^ 30 min later with 0.1 M PBS followed by 4% PFA (in 0.1 M PBS). Spinal cords were dissected and post-fixed in 4% PFA (in 0.1 M PBS) at 4 °C overnight and placed in 0.1 M PBS containing 0.03% sodium azide. Immunolabelling of the samples was performed by first pretreating with methanol in 5 ml Eppendorf tubes by dehydrating with a methanol/H_2_O series at 1 h each at room temperature with shaking at 60 RPM: 20%, 40%, 60%, 80% and 100%. This procedure was followed by 1 h washing with 100% methanol before chilling the samples at 4 °C. Samples were then incubated overnight with shaking in 66% dicholoromethane/33% methanol at room temperature. The samples were washed twice in 100% methanol with shaking at room temperature and then bleached in chilled fresh 5% H_2_O2 in methanol overnight at 4 °C. Samples were rehydrated with a methanol/H_2_O series: 80%, 60%, 40%, 20% and 0.1M PBS, each for 1 h at room temperature under shaking. Samples were washed for 1 h × 2 at room temperature in PTx.2 buffer (0.1 M PBS with 0.2% Triton X-100) under shaking. This was followed by an incubation in 5 ml of permeabilization solution (400 ml PTx.2, 11.5 g glycine, 100 ml DMSO for a total stock volume of 500 ml) for 2 days at 37 °C with shaking at 60 RPM. Samples were incubated in 5 ml of blocking solution (42 ml PTx.2, 3 ml of normal donkey serum, 5 ml of DMSO for a total stock volume of 50 ml) for 2 days at 37 °C with shaking. The samples were incubated for 7 days at 37 °C with shaking in primary antibody solution consisting of PTwH (0.1 M PBS, 2 ml Tween-20, 10 mgl^1^ heparin, 5% dimethyl sulfoxide, 3% normal donkey serum), and cFos antibody (1:2000, Synaptic Systems, 226003) for a total volume of 5 ml per sample. Samples were washed in PTwH for 24 h with shaking and incubated for 7 days at 37 °C with shaking in secondary antibody solution consisting of PTwH, 3% normal donkey serum and donkey anti-rabbit Alexa Fluor 647 (1:400, ThermoFisher Scientific) in a total volume of 5 ml per sample. Samples were washed in PTwH for 24 h with shaking at room temperature. Clearing of the samples was performed by first dehydrating the samples in a methanol/H_2_O series as follows: 20%, 40%, 60%, 80% and 100% twice each for 1 h with shaking at room temperature followed by a 3 h incubation with shaking in 66% dichloromethane/33% methanol at room temperature. Samples were incubated in 100% dichloromethane 15 min twice with shaking to wash residual methanol. Finally, samples were incubated in 100% dibenzyl ether without shaking for refractive index matching of the solution for at least 24 h prior to imaging.

### Tissue clearing (CLARITY)^26,83,86^

We incubated samples in X-CLARITY hydrogel solution (Logos Biosystems Inc., South Korea) for 24 h at 4 °C with gentle shaking. Samples were degassed and polymerized using the X-CLARITY Polymerisation System (Logos Biosystems Inc., South Korea), followed by washes in 0.001 M PBS for 5 minutes at room temperature. Samples were next placed in the X-CLARITY Tissue Clearing System (Logos Biosystems Inc., South Korea), set to 1.5 A, 100 RPM, 37 °C, for 29 h. Clearing solution was made in-house with 4% sodium dodecyl sulfate (SDS), 200mM boric acid with dH_2_O, pH adjusted to 8.5. Following this, samples were washed for at least 24 h at room temperature with gentle shaking in 0.1 M PBS solution containing 0.1% Triton X-100 to remove excess SDS. Finally, samples were incubated in 40 g of Histodenz dissolved in 30 ml of 0.02M PB, pH 7.5, 0.01% sodium azide (refractive index 1.465) for at least 24 h at room temperature with gentle shaking prior to imaging.

### 3D imaging

We performed imaging of cleared tissue using either a customized mesoSPIM^26,87^ and CLARITY-optimized light-sheet microscope (COLM)^26^. A custom-built sample holder was used to secure the central nervous system in a chamber filled with RIMS. Samples were imaged using either a 1.25x or 2.5x objective at the mesoSPIM^26,87^ and a 4x or 10x objective at the COLM^26^ with one or two light sheets illuminating the sample from both the left and right sides. The voxel resolution in the x-, y- and z directions was 5.3 *µ*m x 5.3 *µ*m x 5 *µ*m for the 1.25x acquisition and 2.6 *µ*m x 2.6 *µ*m x 3 *µ*m for the 2.5x acquisition. The voxel resolution of the COLM was 1.4 *µ*m x 1.4 *µ*m x 5 *µ*m for the 4x and 0.59 *µ*m x 0.59 *µ*m x 3 *µ*m for the 10x acquisition. Images were generated as 16-bit TIFF files and then stitched using Arivis Vision4D (Arivis AG, Munich, Germany). 3D reconstructions and optical sections of raw images were generated using Imaris (bitplane, v.9.8.2) software.

### cFos quantifications^27^

For the cleared spinal cords, cFos positive neurons of cleared samples were quantified using Arivis Vision4D (Arivis). After defining a region of interest around the grey matter, each sample was subjected to a custom-made pipeline. We applied morphology, denoising, and normalization filters to enhance the signal of bright objects and homogenized the background. Threshold-based segmentation of the cFos signal was applied within predefined 3D regions to quantify the total number of cFos positive cells. Image analysis parameters were kept constant among all samples. The number of cFos positive cells and their coordinates enabled us to quantify the neuronal activity segment by segment. For the classic immunohistochemistry, the quantification was done on Imaris (bitplane, v.9.8.2) using the spot detection function.

### Axon and synapse quantification^29^

To determine spatial enrichment of axon and synapse density within the grey horn of the spinal cord we implemented a custom image analysis pipeline that includes preprocessing, registration and combination of histological images from different animals. In brief, we implemented all preprocessing in Fiji, and all registration procedures in R, using the image analysis package ‘imageR’, and medical image registration package ‘RNiftyReg’. After dynamic registration, all data were summarized and final quantifications were completed using custom R scripts.

### Neuron-specific recordings and analysis^27^

Spinal cord injury at T4 and infusion of AAV5-Syn-flex-Chrimson was made in the lower thoracic spinal cord of Vsx2^Cre^ mice at least four weeks prior to terminal experiments. Mice were anesthetized with urethane and isoflurane. Two-shank, multi-site electrode arrays (NeuroNexus A2x32-6mm-35-200-177) were lowered into the spinal cord to a depth of 1000 *µ*m, with shanks arranged longitudinally at 350 *µ*m from midline. Signals were recorded with a NeuroNexus Smartbox Pro using a common average reference and while applying 50 Hz notch and 450–5,000 Hz bandpass filters. Stimulation was controlled with a Multi-Channel Systems STG 4004 and MC Stimulus II software. ChrimsonR-expressing neurons were identified using optogenetic stimulation. Twenty pulse trains of 10-ms pulses were delivered at 10 Hz from a 635 nm laser (LaserGlow Technologies LRD-0635-PFR-00100-03). Laser light was delivered to the surface of the spinal cord through a fiber optic cable attached to 400 *µ*m, 0.39 NA cannula with a 5 mm tip (Thorlabs). Optical power was set to 2.35 mW at the tip. Electrical stimulation (EES) consisted of 5 ms pulses delivered every at 1 Hz. EES was delivered with a micro fork probe (Inomed, 45 mm straight, item no. 522610) positioned along the midline just caudal to the recording array. Spike sorting was performed with SpyKING CIRCUS v.1.0.773. The median-based average electrical stimulation artifacts for each channel were subtracted from the recordings prior to sorting. Due to the size and variability of the artifacts, periods containing residual stimulation artifacts were not sorted (–0.5 to +1.5 ms and -0.5 to + 1 ms around stimulus onset for EES and laser stimulation onset, respectively). Sorting results were manually curated using Phy (https://github.com/cortexlab/phy). Single unit clusters were selected for analysis based on their biphasic waveforms and template amplitudes above 50 *µ*V, as well as strong refractory period dips in their spike autocorrelograms. Similar clusters were merged according to the Phy manual clustering guide. ChrimsonR-expressing, putative SC^Hoxa7::Nfib::Vsx2^ neurons were identified based on their low-latency and low-jitter responses to light pulses. Neurons responding to EES or tail pinch were identified by a one-sided Wilcoxon signed-rank test to compare the instantaneous firing rate of units 100 ms before and 100 ms after (EES) or 2 s before and 2 s after (pinch) stimulus onset. For EES, a post-stimulus onset firing rate increase of P value less than 0.001 was used, while for pinch a P value of 0.05 was used due to the necessarily lower number of trials and larger calculation window (minimum 6 trials for pinch, 60 for EES).

### Statistics, power calculations, group sizes and reproducibility

All data are reported as mean values and individual data points. No statistical methods were used to predetermine sample sizes, but our sample sizes are similar to those reported in previous publications^83^. Hemodynamic assays were replicated three to five times, depending on the experiment, and averaged per animal. Statistics were then performed over the mean of animals. All statistical analysis was performed in R using the base package ‘stats’, with primary implementation through the ‘tidyverse’ and ‘broom’ packages. Tests used included one or two-tailed paired or independent samples Student’s t-tests, one-way ANOVA for neuromorphological evaluations with more than two groups, and one- or two-way repeated-measures ANOVA for hemodynamic assessments, when data were distributed normally, tested using a Shapiro–Wilk test. Post hoc Tukey tests were applied when appropriate. For regressions, mixed model linear regression was used in cases of multiple observations, or else standard linear modelling. In cases where group size was equal to or less than three null hypothesis testing was not completed. The significance level was set as P < 0.05. Exclusions of data are noted in the relevant methods sections. Unless stated otherwise, experiments were not randomized, and the investigators were not blinded to allocation during experiments and outcome assessment.

### Single-nucleus RNA sequencing

Single-nucleus dissociation of the mouse lower thoracic and lumbosacral spinal segments was performed according to our established procedures^39,41^. Following euthanasia by isoflurane inhalation and cervical dislocation, the lumbar spinal cord site was immediately dissected and frozen on dry ice. Spinal cords were doused in 500 *µ*l sucrose buffer (0.32 M sucrose, 10 mM HEPES [pH 8.0], 5 mM CaCl_2_, 3 mM Mg acetate, 0.1 mM EDTA, 1 mM DTT) and 0.1% Triton X-100 with the Kontes Dounce Tissue Grinder. 2 ml of sucrose buffer was then added and filtered through a 40-*µ*m cell strainer. The lysate was centrifuged at 3200 g for 10 min at 4 °C. The supernatant was then decanted, and 3 ml of sucrose buffer was added to the pellet for 1 min. We homogenized the pellet using an Ultra-Turrax and 12.5 ml of density buffer (1 M sucrose, 10 mM HEPES [pH 8.0], 3 mM Mg acetate, 1 mM DTT) was added below the nuclei layer. The tube was centrifuged at 3200 g at 4 °C and supernatant poured off. Nuclei on the bottom half of the tube wall were collected with 100 *µ*l PBS with 0.04% BSA and 0.2 U/*µ*l RNase inhibitor. Finally, we resuspended nuclei through a 30 *µ*m strainer, and adjusted to 1000 nuclei/*µ*l.

### Library preparation

snRNA-seq library preparation was carried out using the 10x Genomics Chromium Single Cell Kit Version 3.1. The nuclei suspension was added to the Chromium RT mix to achieve loading numbers of 2000-5000. For downstream cDNA synthesis (13 PCR cycles), library preparation and sequencing, the manufacturer’s instructions were followed.

### Read alignment

We aligned reads to the most recent Ensembl release (GRCm38.93) using Cell Ranger, and obtained a matrix of unique molecular identifier (UMI) counts. Seurat^66^ was used to calculate quality control metrics for each cell barcode, including the number of genes detected, number of UMIs, and proportion of reads aligned to mitochondrial genes. Low-quality cells were filtered by removing cells expressing less than 200 genes or with more than 5% mitochondrial reads. Genes expressed in less than three cells were likewise removed.

### Clustering and integration

Prior to clustering analysis, we first performed batch effect correction and data integration across the two different experimental conditions as previously described^66^. Gene expression data was normalized using regularized negative binomial models^88^, then integrated across batches using the data integration workflow within Seurat. The normalized and integrated gene expression matrices were then subjected to clustering to identify cell types in the integrated dataset, again using the default Seurat workflow. Cell types were manually annotated on the basis of marker gene expression, guided by previous studies of the mouse spinal cord^39,89–91^. Local and projecting neuronal subpopulations were annotated on the basis of *Nfib* and *Zfhx3* expression, respectively^28^. Following our projection-specific snRNA-seq experiment in uninjured mice, each subsequent experiment was re-integrated with this dataset prior to subpopulation annotation. This enabled the identification of the same^28^ neuronal subpopulations across the three distinct experiments^66^.

### Cell type prioritization with Augur

To identify neuronal subpopulations perturbed during natural repair, we implemented our machine-learning method Augur^39,40^. Augur was run with default parameters for all comparisons. To evaluate the robustness of cell type prioritizations to the resolution at which neuronal subtypes were defined in the snRNA-seq data, we applied Augur at various clustering resolutions, and visualized the resulting cell type prioritizations both on a hierarchical clustering tree^92^ of neuron subtypes and as a progression of UMAPs. The key assumption underlying Augur is that cell types undergoing a profound response to a perturbation should become more separable, within the highly multi-dimensional space of gene expression, than less affected cell types. Briefly, Augur withholds a proportion of sample labels, then trains a random forest classifier to predict the condition from which each cell was obtained. The accuracy with which this prediction can be made from single-cell gene expression measurements is then evaluated in cross-validation, and quantified using the area under the receiver operating characteristic curve (AUC).

### Clinical studies design and objectives

All experiments were carried out as part of 3 clinical safety (primary objective) and preliminary efficacy (secondary objectives) trials: **STIMO-HEMO** (NCT04994886, CHUV, Lausanne, Switzerland), **HEMO** (NCT05044923, University of Calgary, Calgary, Canada) and **HemON** (NCT05111093, CHUV, Lausanne, Switzerland). All three trials received approval by the local ethical committees and national competent authorities. All participants signed a written informed consent before their participation. All participants had the option to indicate consent for the publication of identifiable images or videos. All surgical and experimental procedures were performed at the investigational hospital sites (Neurosurgery Department of the Lausanne University Hospital (CHUV) and the Neurosurgery Department of the Foothills Medical Center (Calgary, Canada). The study involved eligibility and baseline assessments before surgery, the surgical implantation of the respective investigational devices, a post-operative period during which EES protocols were configured, and long-term follow-up periods. To date, a total of ten participants have been participating for more than 6 months in the study. More detailed information about these trials can be found in other publications^93^.

### Study participants

Ten individuals who had suffered a traumatic spinal cord injury participated overall in the four studies. Demographic data and neurological status evaluated according to the International Standards for Neurological Classification of Spinal Cord Injury^94^, can be found in other publications^93^.

### Neurosurgical intervention

The participant is put under general anesthesia and is placed in a prone position. Preoperative surgical planning informs the neurosurgeon about the vertebral entry level and predicted optimal position. Based on this knowledge, lateral and anteroposterior fluoroscopy x-rays are performed intraoperatively to guide the location of the laminotomies. A midline skin incision of approximately 5 cm on the back is performed, the fascia opened and the muscles retracted bilaterally. Excision of the midline ligamentous structures and a laminotomy at the desired entry level enables the insertion of the paddle array at the spinal thoracic level. For participants of the STIMO-HEMO and HEMO trials, a second skin incision or extended opening caudally is made and a second laminotomy is performed in the lumbar area based on the pre-operative planning to allow for the insertion of the lumbar paddle lead. The paddle lead(s) (Specify 5-6-5, Medtronic Inc, Minneapolis, MN, USA or ARC^IM^ Thoracic Lead, ONWARD Medical N.V, Eindhoven, Netherlands) are inserted and placed over the midline of the exposed dura-mater and advanced rostrally to the target position guided by repeated fluoroscopies. Electrophysiological recordings are conducted using standard neuromonitoring systems (IOMAX, Cadwell Industries, Kennewick, USA or ISIS Xpress, Inomed Medizintechnik, Emmendingen, Germany). Single-pulses of EES (0.5 Hz) are delivered at increasing amplitude to elicit muscle responses that are recorded from subdermal (Neuroline Twisted Pair Subdermal, 12 x 0.4 mm, Ambu A/S, Ballerup, Denmark) or intramuscular needle electrodes (Inomed SDN electrodes, 40 x 0.45 mm, Inomed Medizintechnik, Emmendingen, Germany) to correct for lateral and rostrocaudal positioning. When the paddles are deviating from a straight midline position, small additional laminotomies are made to remove bony protrusions and guide the paddle to a midline placement. Once the final position is achieved, the leads are anchored to the muscular fascia. In the STIMO-HEMO and HemON trials, the back opening is temporarily closed and the participant is put in lateral decubitus. Subsequently, the back incision is reopened and an abdominal incision of about 5 cm is made per IPG and a subcutaneous pocket is created. In the HEMO trial, incisions of about 5cm are made bilaterally in the upper buttocks region and subcutaneous pockets are created. The paddle array cables are then tunneled between the back opening and subcutaneous pockets to be connected to the IPGs (Intellis, Medtronic Inc, Minneapolis, MN, USA or ARC^IM^ IPG, ONWARD Medical N.V., Eindhoven, Netherlands). The IPGs are implanted in the subcutaneous pockets and all incisions are finally closed.

### Stimulation optimization

Spatial mapping was guided by the pre-clinical mechanisms described previously^19^, and from prior clinical mappings^93^, and was conducted in three steps: 1) intraoperative mapping to identify which rows of electrodes target the *hemodynamic hotspot*^19^, and elicit the largest pressor response in the thoracic spinal cord (T10, T11, T12), 2) post-operative imaging and spinal reconstructions were used to estimate the electrodes that maximize recruitment of the hemodynamic hotspots, 3) a single 2-hour, post-operative mapping session was done to test each row of electrodes on the lead, and pick the three configurations with the largest pressor responses. These configurations were tested in both multipolar and monopolar settings and were validated by personalized simulations to ensure we were optimally targeting the hemodynamic hotspots. Stimulation frequency was defined empirically at 120 Hz for the spatial mapping^19,93^. The pulse width was 300 µs. The amplitude was set by incrementally increasing the current per configuration until the systolic pressure increased by 20 mmHg, the diastolic pressure increased by 10 mmHg, or the patient did not report any discomfort such as muscle contractions, or sensations such as tingling. These mappings were done in a seated position to mimic relevant, daily life orthostatic challenges.

### Hemodynamic monitoring

Beat-to-beat blood pressure and heart rate were obtained via finger plethysmography (Finometer, Finapres Medical Systems; Amsterdam, Netherlands). Beat-by-beat blood pressure was calibrated to brachial artery blood pressure collected using an arm cuff embedded and synchronized with the Finometer^95–98^. Brachial arterial pressure was sampled at 200 Hz, while the systolic, diastolic, and mean arterial pressure were extracted from the calibrated arterial pressure at 1 Hz. The heart rate was also sampled at 1 Hz. Raw data and automatically extracted hemodynamic parameters were saved and exported from the Finometer.

### Orthostatic challenge with tilt table test

Participants were transferred to a supine position on a table capable of head-up tilt. We applied restraint straps to secure the patient below the knees, across the thighs, and the trunk, with the feet stabilized. Resting supine blood pressure was recorded continuously for approximately 5-10 minutes to establish baseline values. Then, we tilted the patient upright up to a maximum of 70 degrees while recording hemodynamic values and symptoms of orthostatic tolerance. The time to reach the desired tilt angle from supine was achieved in less than 45 seconds. Participants were tilted until reaching their tolerance threshold or for a maximum duration of 10 minutes. They were asked not to talk during the test except to inform and grade symptoms. The participant was asked to report any symptoms every 1-3 minutes. The participant was asked to rank their symptoms between 1-10, 1 being no symptoms at all, and 10 being feelings of dizziness, lightheadedness^72^, or nausea^72,99^. The patient was instructed to notify the research team if they needed to be returned to the supine position.

### Postoperative blood pressure data

During a tilt test (See **Orthostatic challenge with tilt table test**), changes in blood pressure were recorded without stimulation or in response to different types of stimulation (continuous or closed-loop stimulation) using the Finometer (see **Post-operative hemodynamic monitoring**). Change in blood pressure or heart rate was defined as the difference in the average of a 60-second window before the start of the tilt and a 20-second window at 3 minutes of the challenge. If the participant could not tolerate at least 3 minutes of the test due to low blood pressure or other symptoms, an average of a 20-second window before the end of the tilt was used. All measurements in seated position were measured with stimulation on for 3 – 5 minutes. Change in blood pressure, or heart rate, was defined as the difference in the average of a 20-second window before the start of EES and the average of a 20-second window at 3 minutes, prior to stopping stimulation. All signals were smoothed over a 10-second window for illustration. The same processing was used for post-operative, day 1 quantification. In the present study, we report on blood pressure data on the 2 study participants implanted with the full ARC^IM^ implantable system.

### Off-label investigational system

The investigational system used in the STIMO-HEMO and HEMO clinical trials consists of a set of CE-marked, FDA-approved medical devices used off-label. Two IPGs (lntellis™ with AdaptiveStim™, Medtronic Inc, Minneapolis, MN, USA) are connected to their respective paddle leads (Specify™ 5-6-5 SureScan™ MRI, Medtronic Inc, Minneapolis, MN, USA), both indicated for chronic pain management. A tablet application with a communicator device (Intellis™ clinician programmer, Medtronic Inc, Minneapolis, MN, USA) is used by the clinical team to wirelessly set up the system and optimize the stimulation parameters. A remote control device and transcutaneous charger device (Patient Programmer and Recharger, Medtronic Inc, Minneapolis, MN, USA) are used by the participants to charge the IPGs and wirelessly turn the stimulation on and off during daily life and adapt a subset of stimulation parameters defined by the clinical team.

### Purpose-built IPG and communication ecosystem for restoring hemodynamic stability

The purpose-built ARC^IM^ IPG developed by ONWARD Medical is a novel 16-channel IPG developed to deliver targeted epidural electrical stimulation. It controls and delivers current-controlled stimulation pulses according to predefined stimulation programs or through commands received in real-time to monopolar or multipolar electrode configurations on 16 channels. The IPG consists of a hermetically sealed, biocompatible can that surrounds the electrical components and a rechargeable battery that enables its function. The IPG is composed of two main components: the header containing the connector block that enables connection with 2 8-contact lead connectors as well as 2 coils for charging and communication, and the can with a rechargeable battery and electronics circuits. The IPG was developed according to all applicable standards for medical device development. Conventional biomedical technologies were used to fabricate the IPG and extensive bench and in-vivo testing was performed to verify its performance. The IPG is implanted subcutaneously at the abdominal level and communicates wirelessly with the ARC^IM^ Hub with Near Field Magnetic Induction (NMFI). This wearable device is worn on a belt over or in proximity to the IPG location and is responsible for wirelessly charging the IPG’s battery and for programming the IPG with stimulation settings received from several user interfaces. The communication between the hub and IPG provides real-time control of stimulation parameters (as fast as 25ms between command and stimulation execution), allowing integration with a fast closed-loop neuromodulation system. The ARC^IM^ Hub contains a Bluetooth Low Energy (BLE) chip to enable fast, reliable wireless communication with external programmers such as the ARC^IM^ Clinician Programmer, an Android app designed for clinicians to configure and test the implanted system and personalized stimulation programs. When a stimulation program is deemed safe for personal use, the Clinician Programmer can be used to make this stimulation program available to the patient. The patients, or their caregivers, can control the system through the ARC^IM^ Personal Programmer. This Android Watch application allows users to select, start, and stop stimulation programs, as well as modulate stimulation amplitudes within predefined safety limits ad-hoc. Device errors, paddle lead impedances, and daily stimulation utilization were extracted from usage logs across all devices. Furthermore, the Clinician Programmer includes an application programming interface (the ARC^IM^ API) that enables other programming softwares to control the stimulation, e.g. for closed-loop control of the stimulation. All devices and softwares are adherent to the applicable standards and their performance was extensively tested. The entire system, including the IPG, received the equivalent of an investigational device exemption from the competent, Swiss authorities.

### Purpose-built paddle lead

The ARC^IM^ Thoracic Lead developed by ONWARD Medical is a new 16-electrode paddle lead that is designed for selective recruitment of the dorsal root entry zones of the low-thoracic spinal cord with optimal coverage of the T10-T12 spinal levels. More detailed information about this paddle lead can be found in other publications^93^.

### Spinal Cord Injury Community Survey (SCICS)

Ethical approval was obtained from an independent ethics board (Veritas Independent Review Board) and also from the Research Ethics Board of Université de Laval (principal investigator’s institution). Ethical approval from local research ethics boards was also obtained to recruit from SCI centers across Canada. Individuals with SCI (n = 1,479) across Canada were recruited using a national consumer awareness campaign and provided written informed consent^76,77^. The survey consisted of a series of variables identified by healthcare and service providers, researchers, as well as individuals with SCI, including demographics, secondary health complications and comorbidities, SCI-related needs, healthcare utilization, community participation, quality of life, as well as overall health ratings^76,77^. Participants were asked how often they had experienced symptoms related to autonomic dysreflexia in the past 12 months and responses were ranked on a 6-point ordinal severity scale ranging from zero (i.e., “Never”) to 5 (i.e., “Every day”). Participants were also asked if they received or sought out treatment in relation to these symptoms on a two-point scale (i.e., “Yes” or “No”), along with the degree to which it limited activities from zero (i.e., “Never”) to 5 (i.e., “Every day”). Participants were also asked if they had experienced specific problems, such as heart disease, in the past 12 months. Participants’ American Spinal Injury Association Impairment Scale (AIS) were estimated using responses to questions about lesion level and sensorimotor/mobility capabilities^76^. A binary approach was used for the evaluation of outcome variables including the level of injury (i.e., cervical SCI vs. non-cervical SCI), the severity of injury (i.e., complete vs. incomplete), presence of autonomic dysreflexia (i.e., yes vs. no), and autonomic dysreflexia symptoms (i.e., yes vs. no). For variables ranked on a 6-point ordinal scale, lower scores (i.e., 0-3) were categorized as “No” and higher scores (i.e., 4-5) were categorized as “Yes”^100^.

### Autonomic Dysfunction following SCI questionnaire (ADF-SCI)

The ADFSCI is a 24-item questionnaire divided into four sections: demographics, medication, autonomic dysreflexia, and orthostatic hypotension. The autonomic dysreflexia section consists of 7 items. Each item employs a 5-point scale to measure the frequency and severity of symptoms related to autonomic dysreflexia, including headaches, goosebumps, heart palpitations, sweating and anxiety, across different situational contexts. Participants were categorized as experiencing symptoms if the item score was higher than 2 or not experiencing symptoms otherwise.

## Acknowledgements

This work was financially supported by: Eurostars project E!113969 PREP2GO; DARPA subaward A21-0795-S001 (P.O. 1083297); Eurostars E!1748 IMPULSE; Medtronic (ERP-2020-12543); Onward Medical, PHRT-279; University of Calgary Reserach Excellence Chair; Brain Canada; Digital Research Alliance of Canada; the Natural Sciences and Engineering Research Council of Canada; the Canadian Institutes of Health Research; Alberta Innovates Health Solutions; Campus Alberta Neuroscience; the Libin Cardiovascular Institute; the Hotchkiss Brain Institute; Hopewell M.I.N.D. Prize; Krembil Research Institute; McCaig Institute for Bone and Joint Health; ONWARD medical; the Swiss National Science Foundation (National Centre of Competence in Research in Robotics, 51NF40185543; Canadian Institutes of Health Research (Graduate Scholarship), Branch Out Neurological Foundation and Eyes High Doctoral Scholarship and NSERC Brain Create to J.E.S.; Ambizione Fellowship to J.W.S., (PZ00P3208988) and subside to G.C. (310030185214 and 310030215668); European Research Council (ERC-2015-CoG HOW2WALKAGAIN 682999; Marie Sklodowska-Curie individual fellowship 842578 to J.W.S.); the Swiss National Supercomputing Center (CSCS); the Swiss National Science Foundation (PZ00P3185728 to M.A.A.); the ALARME Foundation (to G.C and M.A.A.); and Wyss Center for Bio and Neuroengineering and Wings for Life (to M.A.A.). We are grateful to Bernard Schneider and Silvia Arber for providing viral vectors; to Jimmy Ravier and Frederic Merlos for the illustrations; and the Advanced Lightsheet Imaging Center (ALICe) at the Wyss Center for Bio and Neuroengineering, Geneva. This work was supported in part using the resources and services of the Gene Expression Core Facility at the School of Life Sciences of EPFL.

## Author contributions

J.E.S., R.H., J.W.S., A.A.P and G.C. conceptualised and designed experiments; J.E.S., R.H., L.M., M.G., A.L., S.C, C.K, T.H., M.T., M.A.A. and J.W.S performed the rodents experiments; R.C., F.G., S.C., J.R., K.L.K., N.H., A.G., R.D., L.A., A.A.P. and J.B. performed the human experiments; J.E.S., R.H., M.G., Y.Y.T., M.A.S., N.H., A.G., Q.B. and J.W.S. analysed the data. X.K., Y.V., E.M.M., S.L., J.W.S., A.A.P and G.C. designed the hardware and software for neurostimulation. J.E.S., R.H., J.W.S., A.A.P and G.C. wrote the paper, and all the authors contributed to its editing.

## Author information

The authors declare competing financial interests: G.C., A.A.P., J.W.S, J.B., R.D. and S.P.L. hold various patents in relation with the present work. G.C, A.A.P. and R.D. are consultants of ONWARD medical. G.C., A.A.P., J.B. and S.P.L. are minority shareholders of ONWARD, a company with direct relationships with the presented work.

## Data availability

Data that supports the findings and software routines developed for the data analysis will be made available upon reasonable request to the corresponding authors at.

## Additional information

Supplementary is available for this paper at: link. Supplementary information is available for this paper at: link. Correspondence and requests for materials should be addressed to G.C., A.A.P. and J.W.S.

---

**Supplementary Fig. 1.**
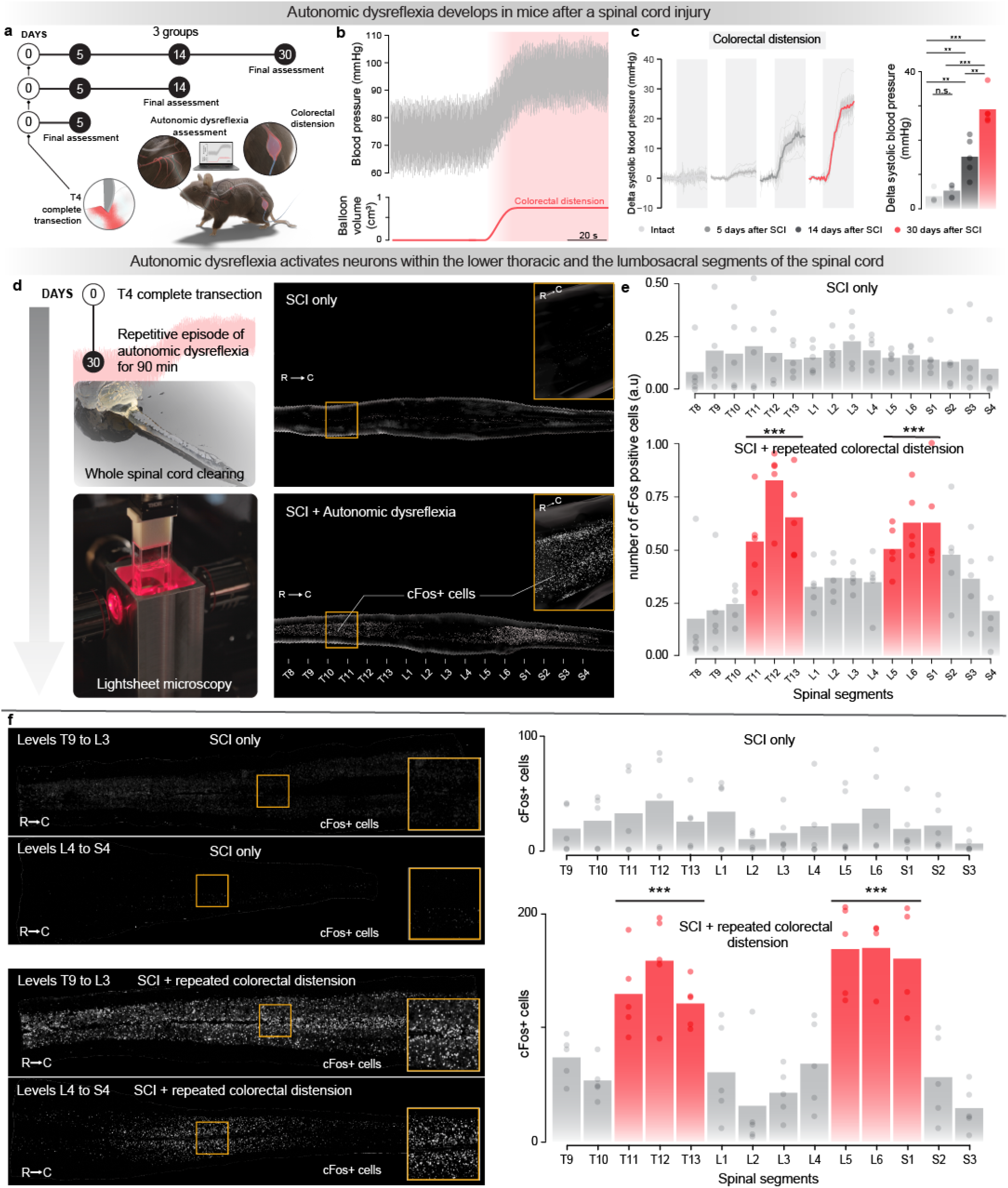
Autonomic dysreflexia triggers transcriptional activity in the lumbosacral and thoracic spinal cord. **a**, Experimental model of autonomic dysreflexia in mice with SCI and timeline of the experiment and final assessments. Blood pressure responses were monitored beat-by-beat using a blood pressure catheter inserted into the carotid artery. Autonomic dysreflexia was elicited using controlled colorectal distension in mice with upper-thoracic SCI. **b**, Changes in blood pressure during autonomic dysreflexia elicited by controlled colorectal distension. **c**,Pressor responses (Left; bold line represents mean trace ± sem for each group and individual line traces are from each animal) and severity of autonomic dysreflexia (Right) measured by the change in systolic blood pressure during colorectal distension at different timepoints after SCI (n = 5 per timepoint). Raw data and statistics provided in **Supplementary Table 1**. **d**, Overview of experimental protocol to identify the regions of the spinal cord activated during autonomic dysreflexia. Thirty days after receiving SCI, mice underwent repetitive episodes of autonomic dysreflexia over 90 minutes, consisting of 30s, and then deflated for 60s. Tissues were collected one hour after the exposure to autonomic dysreflexia, and then processed to visualize the expression of cFos in neurons. *Bottom*, CLARITY-optimized light sheet microscopy of the cleared spinal cord enabled vizualisation of neurons expressing cFos over the entire thoracolumbosacral spinal cord^22,23^ in mice with SCI and mice with SCI that underwent repetitive episodes of autonomic dysreflexia. **e**, Quantification of cFos-labelled neurons in mice with SCI only and mice with SCI that underwent repetitive episodes of autonomic dysreflexia (mixed effect linear model; t = 6.60; p-value = 0.000213). **f**, Quantifications of cFos expression over the whole spinal cord were confirmed with immunohistochemistry and labelling for cFos on longitudinal sections of spinal cord, as illustrated in the representative photomicrographs of spinal cord sections from mice with SCI and mice with SCI that underwent repetitive episodes of autonomic dysreflexia (mixed effect linear model; t = 6.11; p-value = 0.000287).

**Supplementary Fig. 2.**
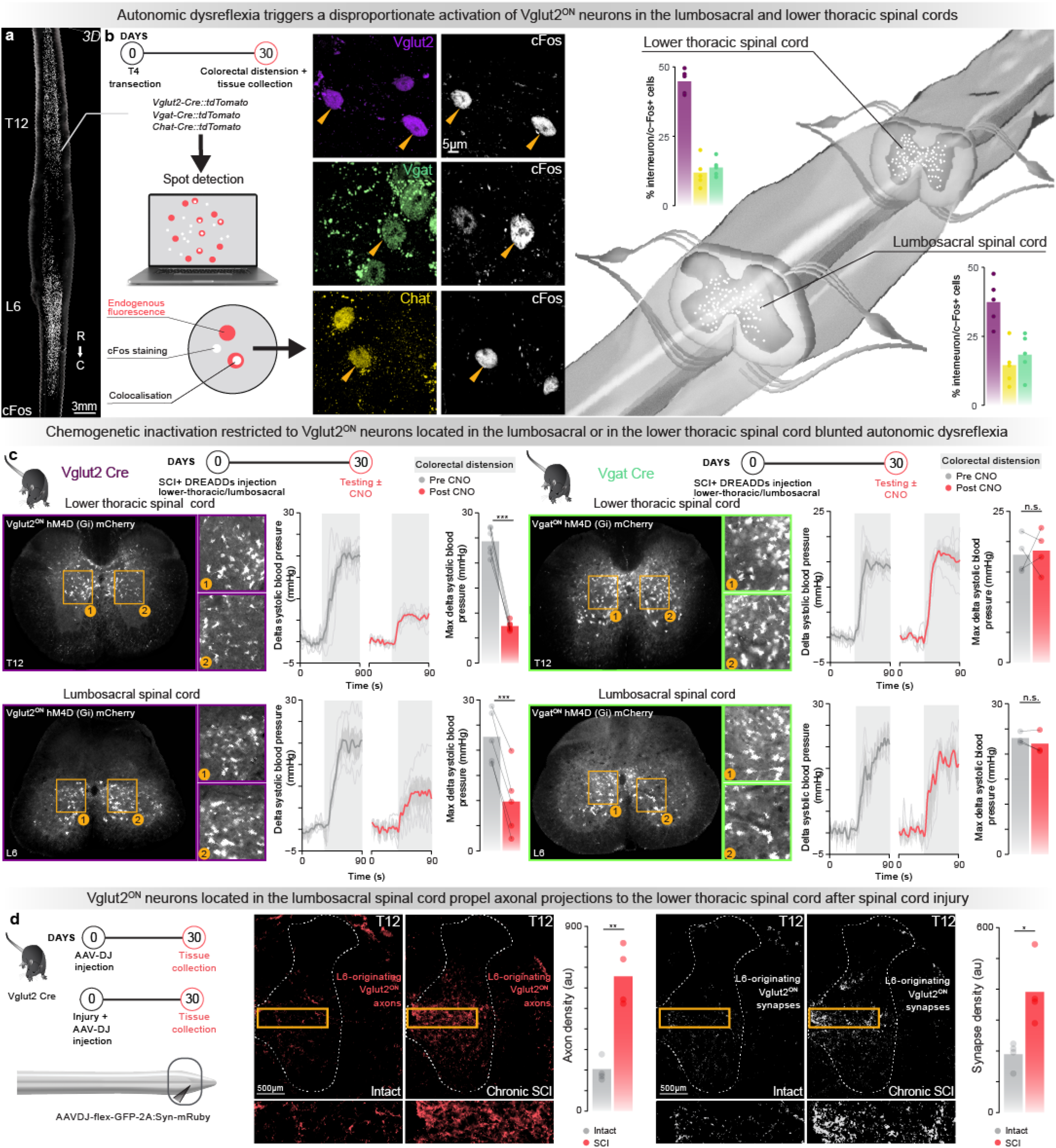
The neurons activated by autonomic dysreflexia. **a**, Schematic overview of experiments to reveal the phenotype of the neurons that are activated during autonomic dysreflexia. We subjected Vglut2^Cre^::Ai9^(RCL-tdT)^, Vgat^Cre^::Ai9^(RCL-tdT)^ and Chat^Cre^::Ai9^(RCL-tdT)^ to repetitive episodes of autonomic dysreflexia at 30 days post-injury. Longitudinal sections of the spinal cord from T9 to L3 and L4 to S4 were immunohistochemically stained for cFos. *Right*, Quantification of colocalisation of cFos-labelled neurons and endogenous fluorescence-tagged neurons (Vglut2^ON^, Vgat^ON^ and Chat^ON^) was performed with automated spot detection (Imaris, Bitplane v.9.8.2). **b**, We next performed experiments to determine the role of these neuronal subpopulations in triggering autonomic dysreflexia. To manipulate the activity of Vglut2^ON^ and Vgat^ON^ neurons, AAV5-hSyn-DIO-hm4D(Gi)-mCherry was infused in either the lower thoracic spinal cord (T11-T13) or lumbosacral spinal cord (L5-S1) of either Vglut2^Cre^ or Vgat^Cre^ mice prior to performing the SCI to express DREADDs in glutamatergic or gabaergic neurons. Photomicrographs of coronal sections from either the lower thoracic or lumbosacral spinal cord in Vglut2^Cre^ and Vgat^Cre^ mice reveal the robust expression of mCherry and thus DREADDs in the targeted neurons. **c**, Chemogenetic inactivation restricted to Vglut2^ON^ neurons located in the lumbosacral (n = 5; paired samples t-test; t = -10.4; p = 0.00048) or lower thoracic spinal cord (n = 5; paired samples t-test; t = -17.5; p = 0.00001) blunted autonomic dysreflexia. In contrast, chemogenetic silencing of Vgat^ON^ neurons in the lumbosacral (n = 3; paired samples t-test; t = -1.53; p = 0.267) or lower thoracic spinal cord (n = 4; paired samples t-test; t = 0.269; p = 0.805) failed to modulate the severity of autonomic dysreflexia. Pressor responses (Left; bold line represents mean trace ± sem for each group and individual line traces are from each animal) and severity of autonomic dysreflexia (bar graph, Right) measured by the change in systolic blood pressure during colorectal distension. **d**. We next aimed to expose the projections from Vglut2^ON^ neurons located in the lumbosacral spinal cord to the Vglut2^ON^ neurons located in the lower thoracic spinal cord. For this, we labeled the axons and synapses of Vglut2^ON^ neurons with infusions of AAV-DJ-hSyn-flex-mGFP-2A-synaptophysin-mRuby into the lumbosacral spinal cord of Vglut2^Cre^. Photomicrographs of the lower thoracic spinal cord from representative mice demonstrate increases in the density of axons (Left; n = 5; independent samples t-test; t = 5.92; p-value = 0.0047) and synaptic puncta (Right; n = 5; independent samples t-test; t = 3.47; p-value = 0.027) emanating from Vglut2^ON^ neurons located in the lumbosacral spinal cord after SCI.

**Supplementary Fig. 3.**
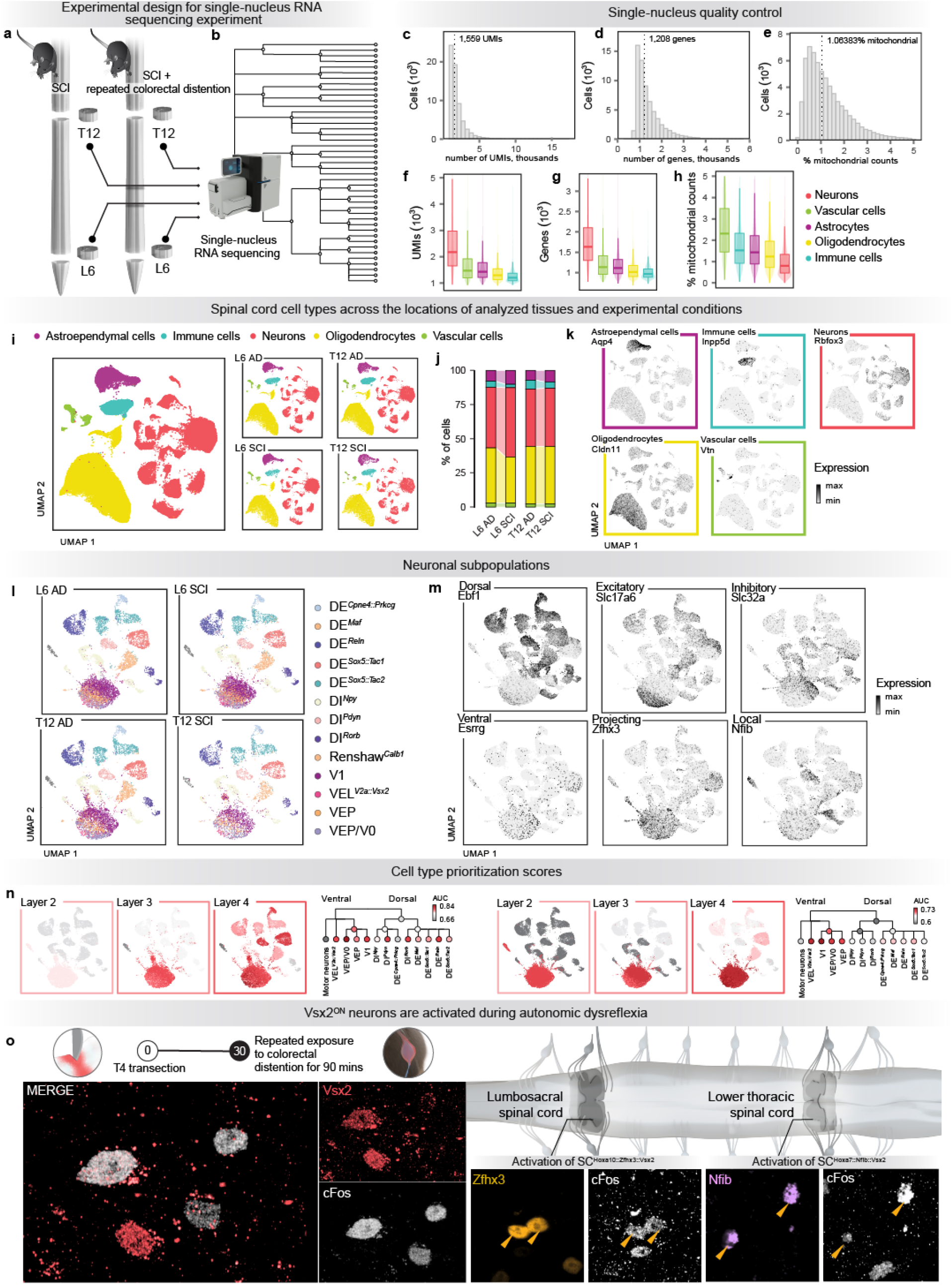
Comparative single-nucleus RNA sequencing atlas of perturbation-responsive neuronal subpopulations during autonomic dysreflexia. **a**, Scheme illustrating our experimental protocol followed by single-nucleus RNA sequencing. Mice received upper-thoracic SCI. After 30 days, half of the mice underwent repetitive episodes of autonomic dysreflexia during 90 minutes. The lumbosacral spinal cord and the lower thoracic were dissected from the mice according to standard procedures. **b**, We obtained high-quality transcriptomes from 64,739 nuclei that were evenly represented across experimental conditions and spatial locations. **c**, Number of unique molecular identifiers (UMIs) per nucleus. Inset text shows the median number of UMIs. **d**, Number of genes detected per nucleus. Inset text shows the median number of genes detected. **e**, Proportion of mitochondrial counts per nucleus. Inset text shows the median proportion of mitochondrial counts. **f**, Number of UMIs quantified per nucleus in each major cell type of the mouse spinal cord. **g**, Number of genes detected per nucleus in each major cell type of the mouse spinal cord. **h**, Proportion of mitochondrial counts per nucleus in each major cell type of the mouse spinal cord. **i**, UMAP visualization of 64,739 nuclei colored by major cell type, segregated by the location of spinal cord tissues (L6, T12) and experimental conditions (SCI only, exposure to repeated episode of autonomic dysreflexia, AD). experimental condition. **j**, Proportions of nuclei from each major cell type depending on the location of spinal cord tissues and experimental conditions. **k**, UMAP visualization showing expression of key marker genes for the major cell types of the mouse spinal cord. **l**, UMAP visualization of 29,144 neuronal nuclei colored by neuronal subpopulations, split by experimental condition. **m**, UMAP visualization showing expression of key marker genes for the major neuronal subpopulation classifications of the mouse spinal cord. **n**, UMAP visualization and dendrograms showing cell type prioritizations assigned by Augur across the neuronal taxonomy of the lower thoracic (*Left*) and lumbosacral (*Right*) spinal cord. **o**, Photomicrographs of the lower thoracic and lumbosacral spinal cord after repetitive episodes of autonomic dysreflexia. Vsx2^ON^ neurons were labelled with immunohistochemistry. Long-distance projecting (Zfhx3, lumbosacral spinal cord) and locally-projecting (Nfib, lower thoracic spinal cord) were additionally colocalized with immunohistochemistry labelling of cFos.

**Supplementary Fig. 4.**
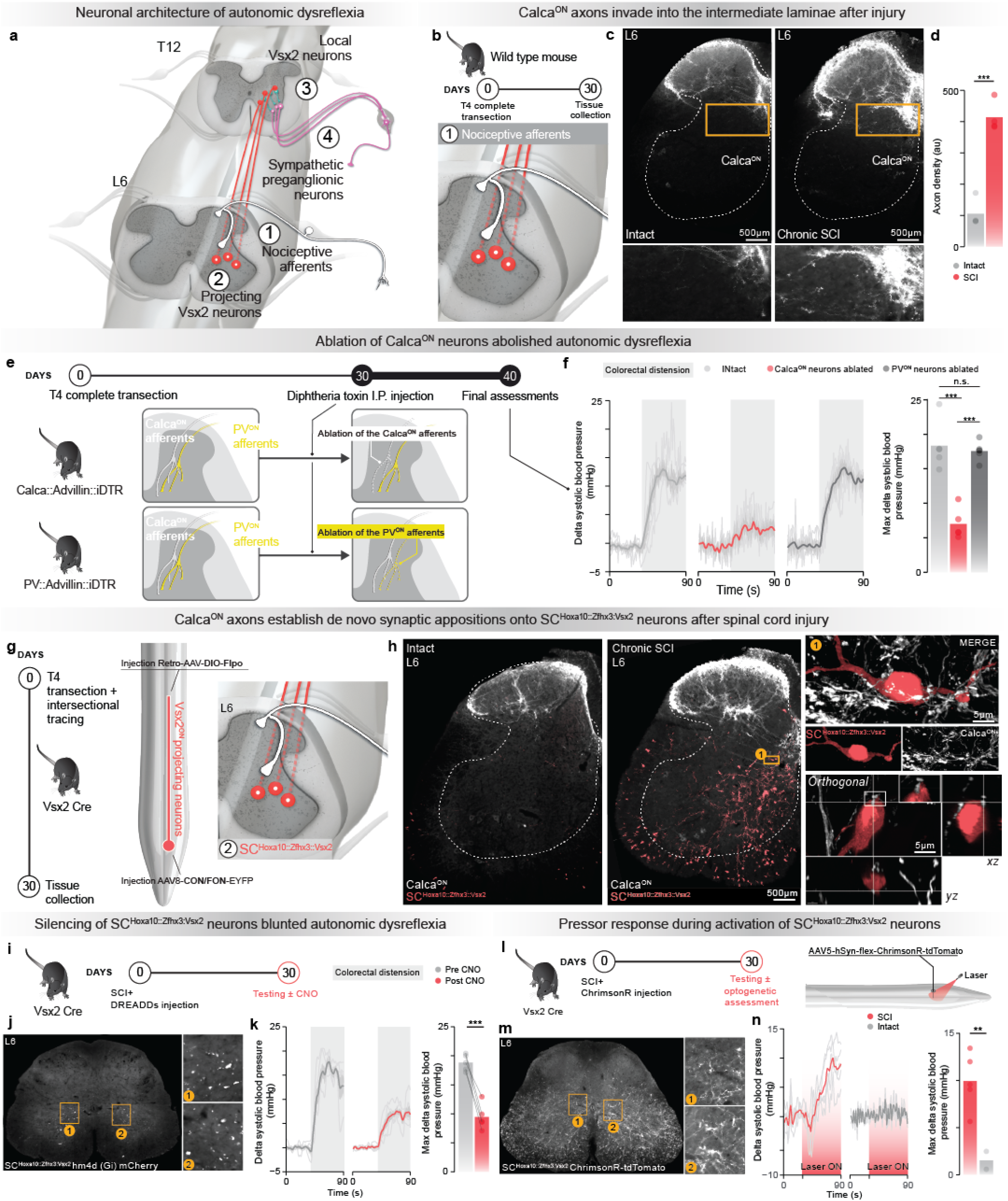
The first and second nodes of the neuronal architecture of autonomic dysreflexia. **a**, Schematic overview of the neuronal architecture of autonomic dysreflexia. **b**, Zoom on the first node of the neuronal architecture of autonomic dysreflexia that involves the growth of projections from Calca^ON^ neurons onto Vsx2^ON^ neurons with long-distance projections, named SC^Hoxa10::Zfhx3::Vsx2^ neurons. This growth was assessed on tissues collected 30 days after SCI in wild-type mice. **c**, Photomicrograph taken at L6 spinal segment from a mouse with an intact spinal cord and a mouse with a chronic SCI in which Calca^ON^ axons were labelled with immunohistochemistry. **d**, Bar plots reporting the density of Calca^ON^ axonal projections into the intermediate laminae of the spinal cord in uninjured mice and mice with chronic SCI (n = 4; independent samples t-test; t = 9.38; p-value = 0.000086). **e**, Overview of the experimental protocol to test the severity of autonomic dysreflexia after the ablation of Calca^ON^ and PV^ON^ neurons. To achieve the ablation of these neurons exclusively in the dorsal root ganglia, we used a Cre- and Flp-dependent strategy in Calca^Cre^::Avil^FlpO^::iDTR and PV^Cre^::Avil^FlpO^::iDTR mice that allowed the expression of diphtheria toxin receptors (DTR) in Calca^ON^ and PV^ON^ neurons located in the dorsal root ganglia, respectively. **f**, Pressor responses (*Left* ; bold line represents mean trace ±standard error of mean (sem) for each group and individual line traces are from each mouse) and severity of autonomic dysreflexia (*Right*) measured by the change in systolic blood pressure during colorectal distension in mice without diphtheria toxin-induced ablation of either Calca^ON^ neurons or PV^ON^ neurons, mice with diphtheria toxin-induced ablation of Calca^ON^ neurons and mice with diphtheria toxin-induced ablation of PV^ON^ neurons (n = 5; independent samples t-test; t= -5.9998; p-value = 0.00064, independent samples t-test; t= -9.3261; p-value = 0.00014). **g**, Zoom on the second node of the neuronal architecture of autonomic dysreflexia that involves SC^Hoxa10::Zfhx3::Vsx2^ neurons projecting to the low thoracic spinal cord. An intersectional viral labelling strategy was used to label the axons of SC^Hoxa10::Zfhx3::Vsx2^ neurons located in the lumbosacral spinal cord and that establish projections in the lower thoracic spinal cord. Vsx2^Cre^ mice received SCI and were injected with Retro-AAV-DIO-FlpO into the lower thoracic spinal cord and AAV8-Con/Fon-EYFP into the lumbosacral spinal cord. **h**, Photomicrograph of the L6 spinal segment from a Vsx2^Cre^ mouse with an intact spinal cord and a Vsx2^Cre^ mouse with a chronic SCI that received intersectional viral tracing to label SC^Hoxa10::Zfhx3::Vsx2^ neurons. Axons from Calca^ON^ were also labelled with immunohistochemistry. Insets show synaptic-like appositions from Calca^ON^ axons onto SC^Hoxa10::Zfhx3::Vsx2^ neurons. **i**, The necessary role of SC^Hoxa10::Zfhx3::Vsx2^ neurons in autonomic dysreflexia was evaluated using Cre-dependent expression of Gi DREADDs in SC^Hoxa10::Zfhx3::Vsx2^ neurons. **j**, Photomicrograph showing the expression of DREADD (G_i_) receptors in SC^Hoxa10::Zfhx3::Vsx2^ neurons. **k**, *Left*, changes in systolic blood pressure in response to colorectal distension (shared area). Bold line represents mean trace ±sem for each group and individual line traces are from each mouse) and severity of autonomic dysreflexia. *Right*, Severity of autonomic dysreflexia in Vsx2^Cre^ mice before and after chemogenetic silencing of Vsx2^ON^ neurons located in the lumbosacral spinal cord (n = 5; paired samples t-test; t = -9.47; p-value = 0.00069). **l**, The sufficient role of SC^Hoxa10::Zfhx3::Vsx2^ neurons in triggering autonomic dysreflexia was evaluated using optogenetic activation of SC^Hoxa10::Zfhx3::Vsx2^ neurons in Vsx2^Cre^ mice injected with AAV-Syn-flex-ChrimsonR-tdTomato into at the lumbosacral spinal cord. 30 days after SCI, blood pressure responses were monitored beat-by-beat using a blood pressure catheter inserted into the carotid artery. Red-shifted light was shined over the lumbosacral spinal cord for 60 seconds during each trial. **m**, Photomicrograph showing the expression of ChrimsonR in SC^Hoxa10::Zfhx3::Vsx2^ neurons. **n**, Left, Changes in systolic blood pressure in response to the photostimulation of SC^Hoxa10::Zfhx3::Vsx2^ neurons in mice with intact spinal cord and with chronic SCI. Bold line represents mean trace ±sem for each group and individual line traces are from each mouse) and blood pressure responses due to optogenetic activation of SC^Hoxa10::Zfhx3::Vsx2^ neurons. *Right*, Bar plots reporting mean changes in blood pressure in Vsx2^Cre^ mice with intact spinal cord and with SCI during optogenetic activation of SC^Hoxa10::Zfhx3::Vsx2^ neurons (n = 5; independent samples t-test; t = 5.14; p-value = 0.00496).

**Supplementary Fig. 5.**
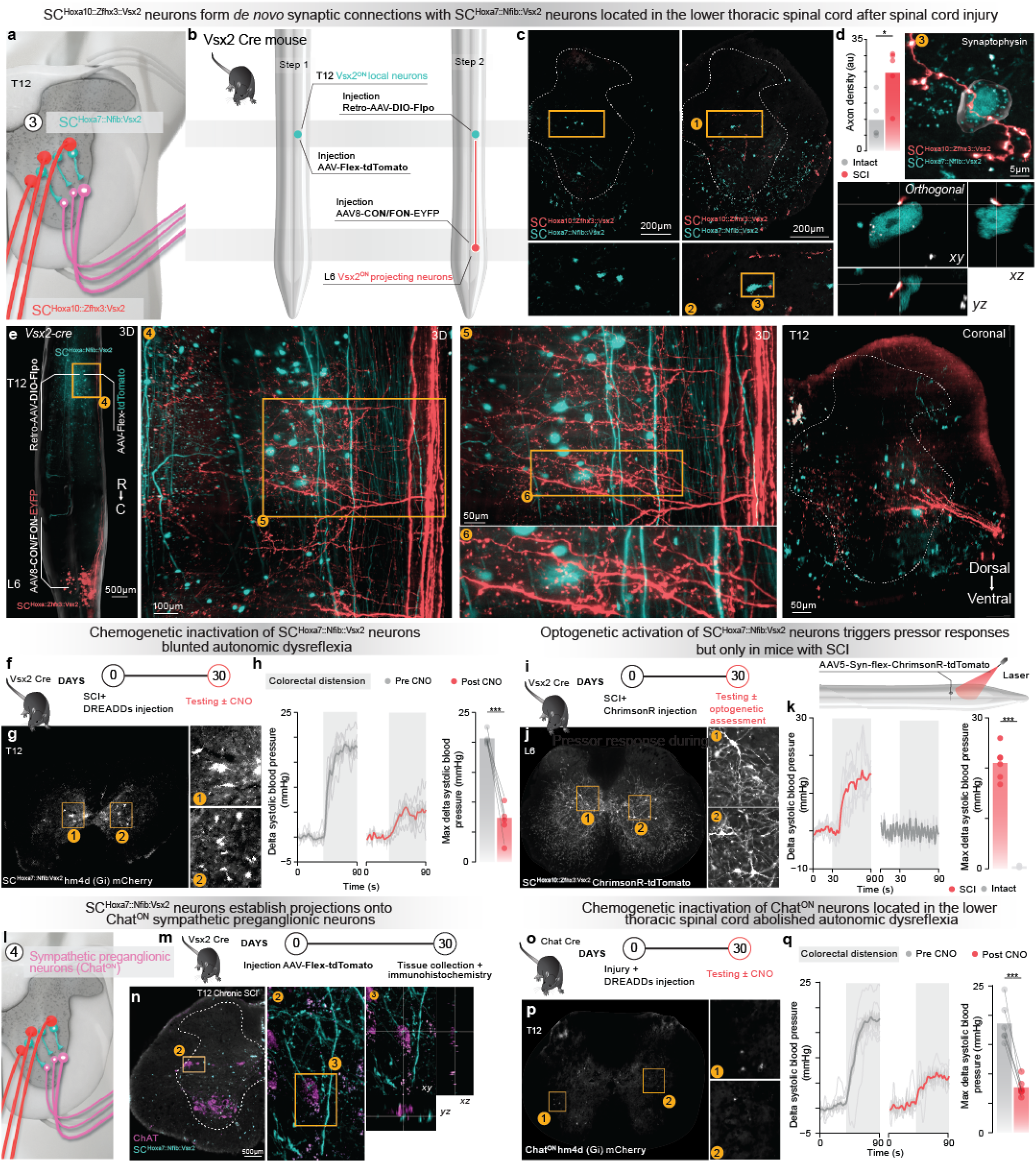
The third and fourth nodes of the neuronal architecture of autonomic dysreflexia. **a**, Zoom on the third node of the neuronal architecture of autonomic dysreflexia that involves SC^Hoxa7::Nfib::Vsx2^ neurons located in the lower thoracic spinal cord. **b**, Overview of intersectional viral tracing strategy to label projections from SC^Hoxa10::Zfhx3::Vsx2^ into the lower thoracic spinal cord concomitantly to the labelling of SC^Hoxa10::Zfhx3::Vsx2^ *Step 1*, AAV5-hSyn-flex-tdTomato was infused into the lower thoracic spinal cord of Vsx2-Cre mice to label SC^Hoxa7::Nfib::Vsx2^. *Step 2*, Retro-AAV-DIO-FlpO was infused into the lower thoracic spinal cord and AAV8-Con/Fon-EYP into the lumbosacral spinal cord to label the projections from SC^Hoxa10::Zfhx3::Vsx2^ neurons located in the lumbosacral spinal cord and that project in the lower thoracic spinal cord. **c**, Photomicrographs of the lower thoracic spinal cord with intersectional viral tracing labelling projections from SC^Hoxa10::Zfhx3::Vsx2^, SC^Hoxa7::Nfib::Vsx2^ neurons and their projections from a representative mouse with an intact spinal cord and mouse with SCI. **d**, Bar plots reporting the mean density of projections from SC^Hoxa10::Zfhx3::Vsx2^ neurons in the grey matter of the lower thoracic spinal cord in mice with an intact spinal cord and with chronic SCI (n = 5; independent samples t-test; t = -3.09; p-value = 0.0162). **e**, Whole spinal cord visualization of projections from SC^Hoxa10::Zfhx3::Vsx2^ neurons located in the lumbosacral spinal cord (red) and visualization of SC^Hoxa7::Nfib::Vsx2^ neurons (blue) located in the lower thoracic spinal cord in mice with chronic SCI. **f**, The necessary role of SC^Hoxa7::Nfib::Vsx2^ neurons in autonomic dysreflexia was evaluated using Cre-dependent expression of G_i_ DREADDs in SC^Hoxa7::Nfib::Vsx2^ neurons. **g**, Photomicrograph showing the expression of G_i_ DREADD receptors in SC^Hoxa7::Nfib::Vsx2^ neurons. **h**, *Left*, changes in systolic blood pressure in response to colorectal distension. Bold line represents mean trace ±sem for each group and individual line traces are from each mouse) and severity of autonomic dysreflexia. *Right*, Severity of autonomic dysreflexia in Vsx2^Cre^ mice before and after chemogenetic silencing of Vsx2^ON^ neurons located in the lower thoracic spinal cord (n = 5; paired samples t-test; t = -9.39; p-value = 0.00072). **i**, The sufficient role of SC^Hoxa7::Nfib::Vsx2^ neurons in autonomic dysreflexia was evaluated using optogenetic activation of SC^Hoxa7::Nfib::Vsx2^ neurons in Vsx2^Cre^ mice injected with AAV-Syn-flex-ChrimsonR-tdTomato into the lower thoracic spinal cord. 30 days after SCI, blood pressure responses were monitored beat-by-beat using a blood pressure catheter inserted into the carotid artery. Red-shifted light was shine over the lumbosacral spinal cord for 60 seconds during each trial. **j**, Photomicrograph showing the expression of ChrimsonR in SC^Hoxa7::Nfib::Vsx2^ neurons. **k**, *Left*, changes in systolic blood pressure in response to colorectal distension. Bold line represents mean trace ±sem for each group and individual line traces are from each mouse) and blood pressure responses due to optogenetic activation of SC^Hoxa7::Nfib::Vsx2^ neurons. *Right*, Blood pressure responses in Vsx2^Cre^ mice with intact spinal cord and with chronic SCI during optogenetic activation of SC^Hoxa7::Nfib::Vsx2^ neurons (n = 5; independent samples t-test; t = 15.4; p-value = 0.0000148). **l**, Zoom on the fourth node of the neuronal architecture of autonomic dysreflexia that involves Chat^ON^ sympathetic preganglionic neurons. **m**, Overview of experimental protocol to label projections from SC^Hoxa7::Nfib::Vsx2^ neurons located in the lower thoracic spinal cord in Vsx2^Cre^ mice with SCI. Thirty days after SCI and viral tracing, the spinal cord tissues were collected and processed. **n**, Photomicrograph of the lower thoracic spinal cord from a mouse with an intact spinal cord and a mouse with chronic SCI in which the projections of SC^Hoxa7::Nfib::Vsx2^ were labelled concomitantly to the immunohistochemical labelling of Chat^ON^ neurons. **o**, The necessary role of Chat^ON^ neurons in autonomic dysreflexia was evaluated using Cre-dependent expression of G_i_ DREADDs in Chat^ON^ neurons. **p**, Photomicrograph illustrating the expression of G_i_ DREADD receptors in SC^Hoxa7::Nfib::Vsx2^ neurons. **q**, As in **h**, for Chat^ON^ neurons located in the lower thoracic spinal cord (n = 5; paired samples t-test; t = -8.03; p-value = 0.00048).

**Supplementary Fig. 6.**
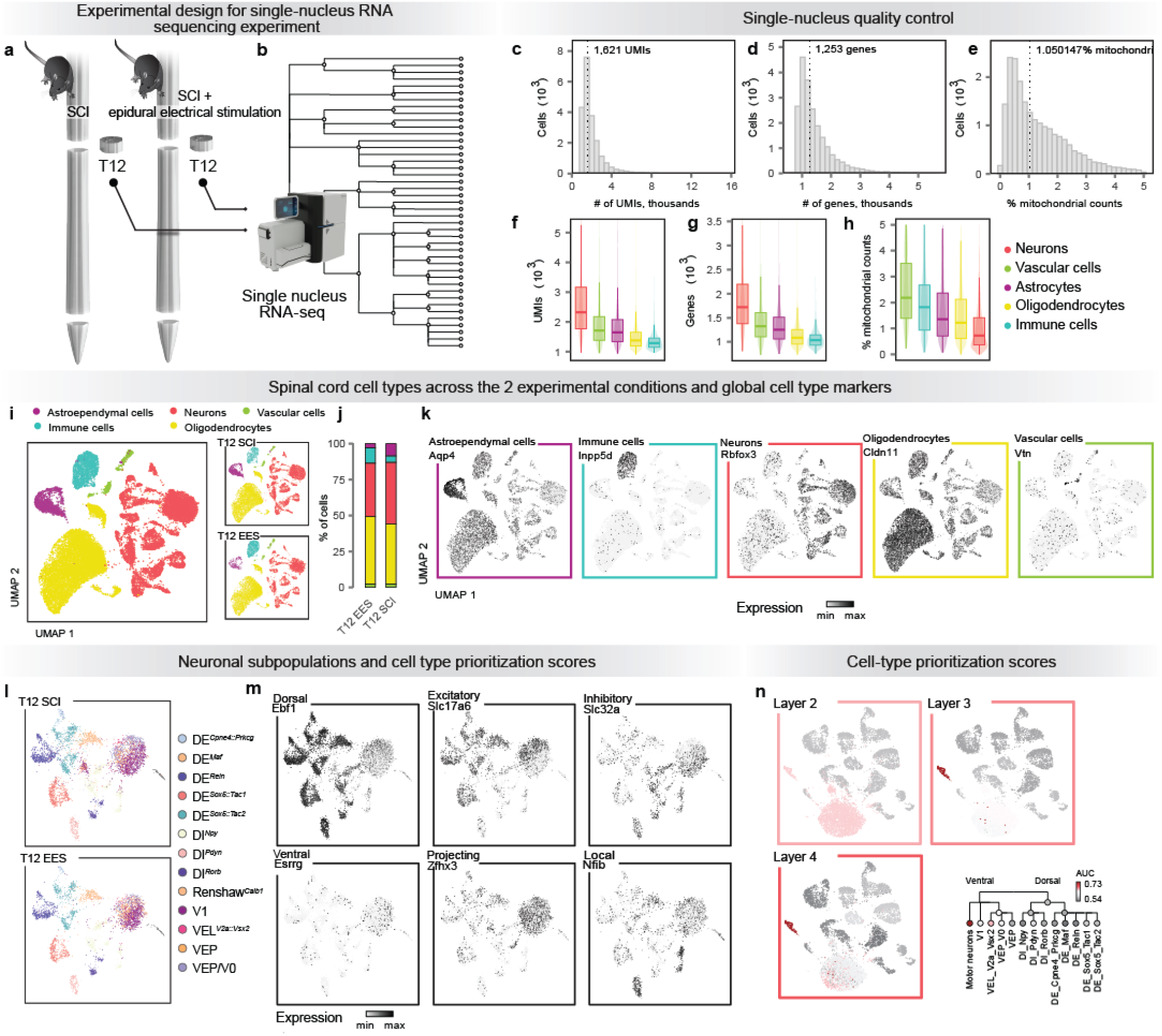
Comparative single-nucleus RNA sequencing atlas of perturbation-responsive neuronal subpopulations during epidural electrical stimulation. **a**, Scheme illustrating the experimental protocol followed by single-nucleus RNA sequencing. Days after SCI for half of the mice, EES was applied continuously over the lower thoracic spinal cord during 45 minutes. The lower thoracic spinal cord was harvested from the mice according to standard procedures. **b**, We obtained high-quality transcriptomes from 21,098 nuclei that were evenly represented across experimental conditions and spatial locations. **c**, Number of unique molecular identifiers (UMIs) per nucleus. Inset text shows the median number of UMIs. **d**, Number of genes detected per nucleus. Inset text shows the median number of genes detected. **e**, Proportion of mitochondrial counts per nucleus. Inset text shows the median proportion of mitochondrial counts. **f**, Number of UMIs quantified per nucleus in each major cell type of the mouse spinal cord. **g**, Number of genes detected per nucleus in each major cell type of the mouse spinal cord. **h**, Proportion of mitochondrial counts per nucleus in each major cell type of the mouse spinal cord. **i**, UMAP visualization of 21,098 nuclei colored by major cell type, split by experimental condition. **j**, Proportions of nuclei from each major cell type across all experimental conditions. **k**, UMAP visualization showing expression of key marker genes for the major cell types of the mouse spinal cord. **l**, UMAP visualization of 8,471 neuronal nuclei colored by neuronal subpopulations, split by experimental condition. **m**, UMAP visualization showing expression of key marker genes for the major neuronal subpopulation classifications of the mouse spinal cord. **n**, UMAP visualization and dendrograms showing cell type prioritizations assigned by Augur across the neuronal taxonomy of the lower thoracic spinal cord.

**Supplementary Fig. 7.**
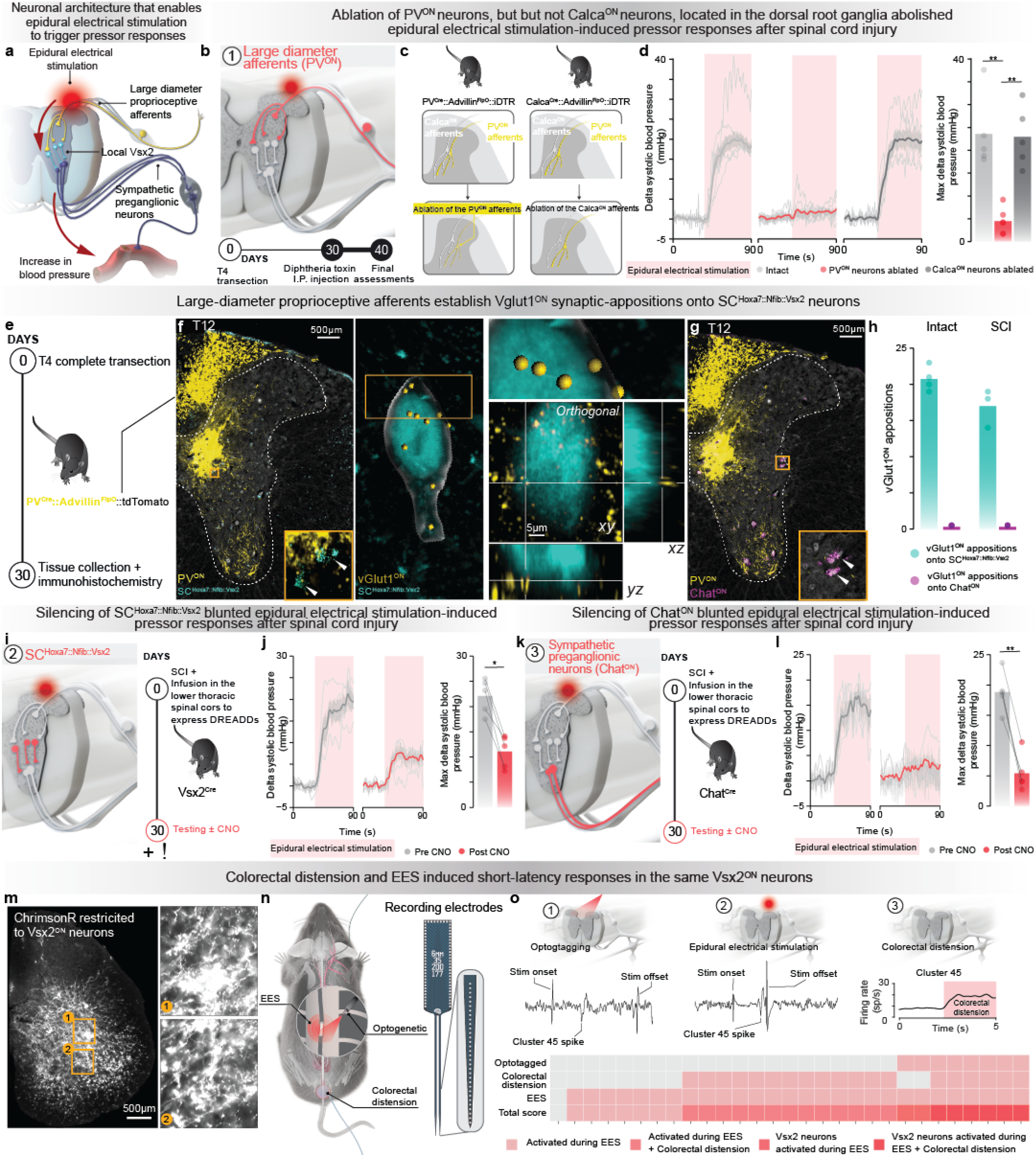
The neuronal architecture activated by EES to induce pressor response. **a**, Schematic overview of the neuronal architecture through which EES induces pressor responses. **b**, Zoom on the first node of the neuronal architecture of EES-induced pressor responses that involves PV^ON^. **c**, Overview of the experimental protocol to test the involvement of afferent fibers from PV^ON^ and Calca^ON^ neurons in EES-induced pressor responses. To achieve the ablation of these neurons exclusively in the dorsal root ganglia, we used a Cre- and Flp-dependent strategy in Calca^Cre^::Avil^FlpO^::iDTR and PV^Cre^::Avil^FlpO^::iDTR mice that allowed the expression of diphtheria toxin receptors (DTR) in these specific neurons. **d**, EES-induced pressor responses (*Left* ; bold line represents mean trace ±sem for each group and individual line traces are from each mouse) (*Right*) measured by the change in systolic blood pressure during EES in mice without any ablation, mice with diphtheria toxin-induced ablation of PV^ON^ neurons and mice with diphtheria toxin-induced ablation of Calca^ON^ neurons (n = 5; independent samples t-test; t= -5.4141; p-value = 0.0043, independent samples t-test; t= 6.3166; p-value = 0.0020). **e**, Overview of the experiment strategy to visualize large-diameter PV^ON^ fibers in PV^Cre^::Avil^FlpO^::Ai9^(RCL-tdT)^ mice and confirmed that they established vGlut1^ON^ synaptic-appositions onto SC^Hoxa7::Nfib::Vsx2^ neurons. Thirty days after SCI, spinal cord tissues were collected and processed. **f**, Photomicrograph of the lower thoracic spinal cord showing vGlut1 synaptic puncta and synaptic-like appositions from large-diameter afferent neurons (PV^Cre^::Advil^FlpO^::Ai9^(RCL-tdT)^ mice) onto SC^Hoxa7::Nfib::Vsx2^ neurons labelled with in situ hybridization (*Left*) or viral tract tracing (*Right*). **g**, Photomicrograph of the lower thoracic spinal cord from a PV^Cre^::Advil^FlpO^::Ai9^(RCL-tdT)^ mouse combined with immunohistochemical labelling of ChatON neurons. **h**, Quantification of vGlut1^ON^ synaptic-appositions onto SC^Hoxa7::Nfib::Vsx2^ neurons and ChatON neurons in PV^Cre^::Advil^FlpO^::Ai9^(RCL-tdT)^ mice with an intact spinal cord and with a chronic SCI. **i**, Zoom on the second node of the neuronal architecture of EES-induced pressor responses that involves SC^Hoxa7::Nfib::Vsx2^. The necessary role of SC^Hoxa7::Nfib::Vsx2^ neurons in EES-induced pressor response was evaluated using Cre-dependent expression of Gi DREADDs in SC^Hoxa7::Nfib::Vsx2^ neurons. **j**, EES-induced pressor responses (*Left* ; bold line represents mean trace ±sem for each group and individual line traces are from each mouse) (*Right*) measured by the change in systolic blood pressure during EES in the same mice before and after chemogenetic silencing of Vsx2^ON^ neurons located in the lower thoracic spinal cord (n = 5; paired samples t-test; t = -4.21; p-value = 0.014). **k**, Zoom on the third node of the neuronal architecture of EES-induced pressor responses that involves Chat^ON^ sympathetic preganglionic neurons. As in **h**, for Chat^ON^ neurons located in the lower thoracic spinal cord. **l**, As in **j**, for Chat^ON^ neurons in the lower thoracic spinal cord (n = 5; paired samples t-test; t = -7.07; p-value = 0.0021). **m**, Photomicrograph showing the expression of ChrimsonR in Vsx2^ON^ neurons and the tract resulting from the insertion of one electrode shank. **n**, Schematic overview of experiments to record the activity of SC^Hoxa7::Nfib::Vsx2^ during the application of EES and during episodes of autonomic dysre-flexia. **o**, *Top*, the waveforms display spikes and firing rate evoked by optogenetic stimulation of Vsx2^ON^ neurons by the application of continuous EES over the lower thoracic spinal cord, and by colorectal distention. Heatmap of neuronal clusters activated by EES, activated by EES and colorectal distension, activated by EES and tagged as Vsx2^ON^ neurons by optogenetic stimulation and EES, and activated by EES and colorectal distension and tagged as Vsx2^ON^ neurons activated by optogenetic stimulation.

**Supplementary Fig. 8.**
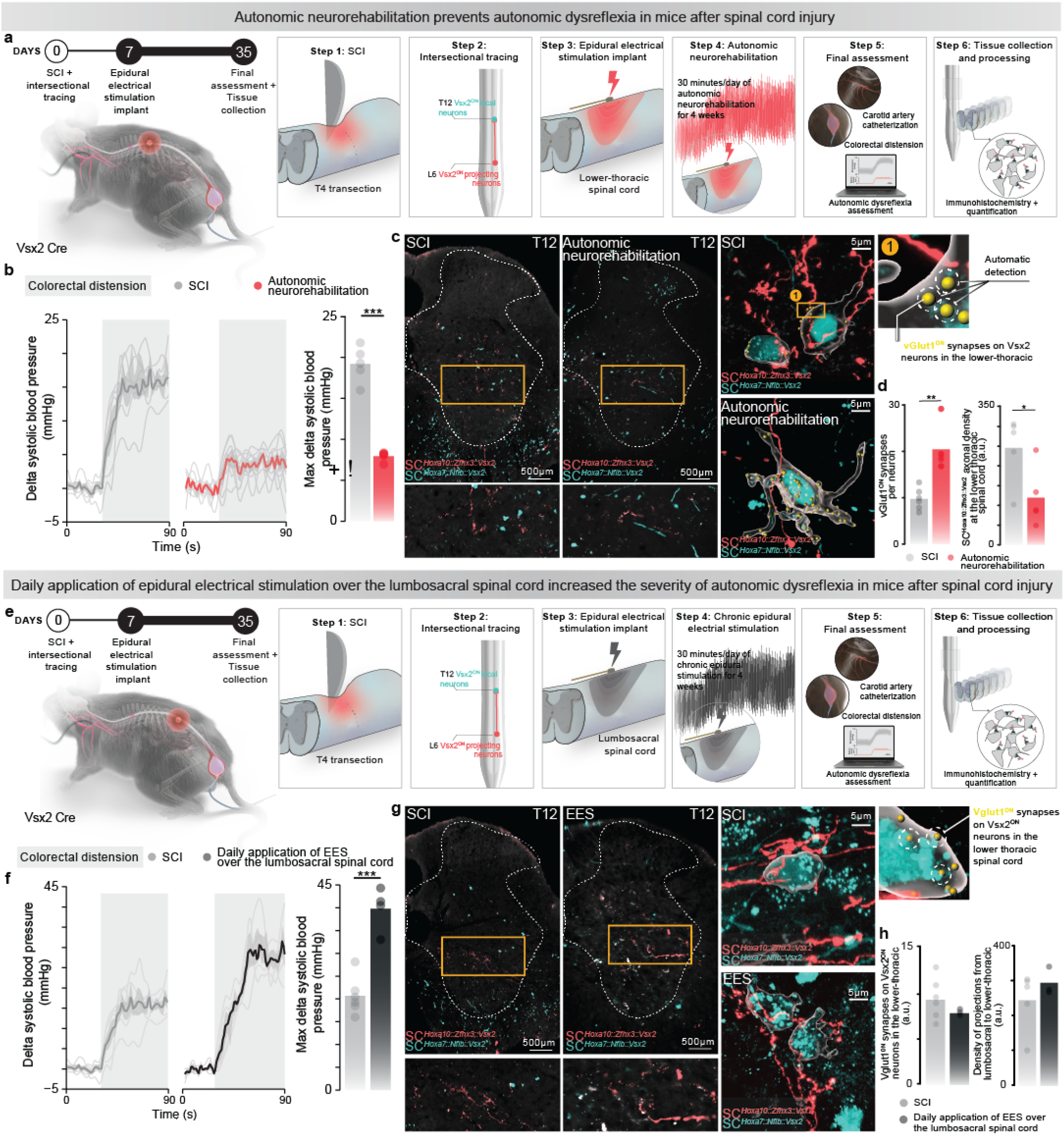
Autonomic neurorehabilitation reversed autonomic dyresflexia in mice with SCI. **a**, Overview of the experimental protocol to deliver autonomic neurorehabilitation in mice with SCI. *Step 1*. Mice received a complete transection of the spinal cord at the level of the T4 segment. *Step 2*.Intersectional viral tracing by infusing Retro-AAV-DIO-FlpO into the lower thoracic spinal cord and AAV8-Con/Fon-EYP into the lumbosacral spinal cord to label SC^Hoxa10::Zfhx3::Vsx2^ neurons located in the lumbosacral spinal cord that project onto SC^Hoxa7::Nfib::Vsx2^ neurons located in the lower thoracic spinal cord. *Step 3*. One week after SCI, electrodes were implanted over the T12 spinal segment to deliver EES. *Step 4*. EES was applied for 30 minutes everyday for 4 weeks. *Step 5*. F of autonomic dysreflexia was assessed during terminal experiments conducted in mice with chronic SCI and mice with chronic SCI that underwent autonomic neurorehabilitation. *Step 6*. Spinal cord tissues were collected and processed. **b**, Changes in systolic blood pressure (*Left* ; bold line represents mean trace ± sem for each group and individual line traces are from each animal) and severity of autonomic dysreflexia (*Right*) measured by the change in systolic blood pressure during colorectal distension in mice with and without autonomic neurorehabilitation (n = 5; independent samples t-test; t = -7.45; p-value = 0.00056). **c**, *Left*, Photomicrographs of the lower thoracic spinal cord in mice with chronic SCI and mice with chronic that underwent autonomic neurorehabilitation in which SC^Hoxa10::Zfhx3::Vsx2^ neurons located in the lumbosacral spinal cord were labelled with an intersection virus strategy concomitantly to the labelling of SC^Hoxa7::Nfib::Vsx2^ neurons. (Right) Photomicrographs of the lower thoracic spinal cord with intersectional viral labelling combined with immunohistochemical labelling of vGlut1^ON^ synapses in mice with chronic SCI and mice with chronic SCI that underwent autonomic neurorehabilitation. vGlut1ON synaptic puncta and synaptic-like appositions from SC^Hoxa10::Zfhx3::Vsx2^ neurons onto SC^Hoxa7::Nfib::Vsx2^ neurons in mice with chronic SCI and mice with chronic SCI that underwent autonomic neurorehabilitation. **d**, *Left*, Bar plots reporting the mean number of vGlut1^ON^ synaptic puncta apposing SC^Hoxa7::Nfib::Vsx2^ neurons (n = 5; independent samples t-test; t = 4.44; p-value = 0.0055), *right*, and the mean density of axonal projections from SC^Hoxa10::Zfhx3::Vsx2^ neurons in the grey matter of the lower thoracic spinal cord in mice with chronic SCI and mice with chronic SCI that underwent autonomic neurorehabilitation (n = 5; independent samples t-test; t = 2.51; p-value = 0.0369). **e**, As in **a**, for mice subjected to daily application of EES over the lumbosacral spinal cord. **f**, As in **b**, for mice subjected to daily application of EES over the lumbosacral spinal cord (n = 5; independent samples t-test; t = 5.82; p-value = 0.00070). **g**, As in **c**, for mice subjected to daily application of EES over the lumbosacral spinal cord. **h**, As in **d**, for mice subjected to daily application of EES over the lumbosacral spinal cord.

**Supplementary Fig. 9.**
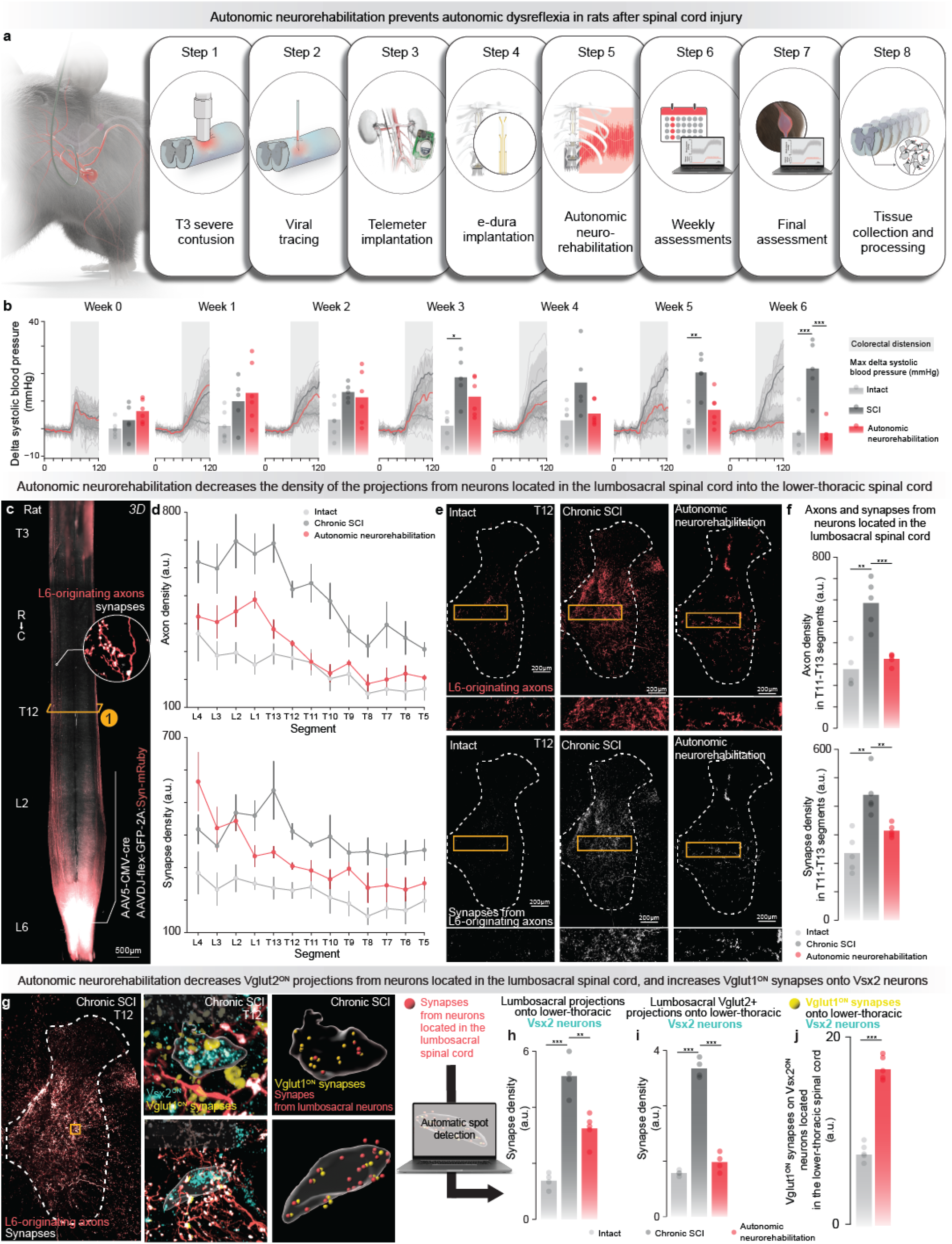
Autonomic neurorehabilitation reversed autonomic dyresflexia in rats with contusion SCI. **a**, Overview of the experimental protocol to deliver autonomic neurorehabilitation in rats with SCI. **Step 1**. Rats received a severe contusion (380 Kdyn) of the spinal cord at the level of T3 segment. **Step 2**. AAV-DJ-hSyn-flex-mGFP-2A-Synaptophysin-mRuby and an AAV-Cre were co-infused into the the L6 segment of the spinal cord to label the projections from neurons located in the lumbosacral spinal cord. **Step 3**. A wireless telemeter recording system, including a blood pressure cannula inserted into the abdominal aorta and microelectrodes sutured over the sympathetic renal nerve, was implanted chronically to monitor hemodynamics and sympathetic nerve activity, respectively. **Step 4**. Seven days after SCI, an an electronic dura mater (e-dura) designed to target the dorsal roots projecting to the T11, T12, and T13 spinal segments was implanted over the hemodynamic hotspot to regulate blood pressure. **Step 5**. EES was applied for 30 minutes everyday during 6 weeks using a proportional-integral (PI) controller that adjusted the amplitude of EES in closed-loop s to augment the systolic blood pressure to a target range. **Step 6**. The severity of autonomic dysreflexia, induced by colorectal distension, was assessed every week for 6 weeks. **Step 7**. After 6 weeks of autonomic neurorehabilitation, a final assessment was performed to test the severity of autonomic dysreflexia in all groups, which included rats with intact spinal cord, rats with chronic SCI and rats with chronic SCI that underwent autonomic neurorehabilitation. **Step 8**. Spinal cords were collected and processed. **b**, Changes in systolic blood pressure in response to colorectal distension (*Left* ; bold line represents mean trace ± sem for each group and individual line traces are from each rat) and bar plots reporting the severity of autonomic dysreflexia (*Right*) measured by the change in systolic blood pressure during colorectal distension over the course of 6 weeks in rats with intact spinal cord, rats with chronic SCI and rats with chronic SCI that underwent autonomic neurorehabilitation. Raw data and statistics provided in **Supplementary Table 1**. **c**, Whole spinal cord visualization of projections from neurons located in the lumbosacral spinal cord. **d**, Plots reporting density of axonal projections (*top*) and synaptic punta (*bottom*) from neurons located in the lumbosacral spinal cord into the grey matter of the lower thoracic spinal cord in rats with intact spinal cord, rats with chronic SCI and rats with chronic SCI that underwent autonomic neurorehabilitation. **e**, Micrographs of the lower thoracic spinal cord in which the axonal projections and synaptic puncta from neurons located in the lumbosacral spinal cord are labelled for the three groups of rats. **f**, Bar plots reporting the mean density of axonal projections and synaptic puncta from neurons located in lumbosacral spinal cord into the grey matter of the lower thoracic spinal cord for the three groups of rats. Raw data and statistics are provided in **Supplementary Table 1**. **g**, Micrographs of the lower thoracic spinal cord in which axonal projections and synaptic puncta from neurons located in the lumbosacral spinal cord are labelled concomitantly to vGlut1^ON^ synapses from large-diameter afferents and Vsx2^ON^ neurons. The density of vGlut1^ON^ synapses onto Vsx2^ON^ neurons is reconstructed for a rat with chronic SCI and a rat with chronic SCI that underwent autonomic neurorehabilitation. **h**, Bar plots reporting the density of synaptic-like appositions from neurons located in the lumbosacral spinal cord onto Vsx2^ON^ neurons in rats with intact spinal cord, rats with chronic SCI, and rats with chronic SCI that underwent autonomic neurorehabilitation. Raw data and statistics are provided in **Supplementary Table 1**. **i**, As in i, for vGlut2^ON^ synaptic puncta onto SC^Hoxa7::Nfib::Vsx2^ neurons. Raw data and statistic provided are in **Supplementary Table 1**. **j**, Quantification of vGlut1^ON^ synaptic puncta from large-diamter afferents Vsx2^ON^ in rats with chronic SCI and rats with chronic SCI that underwent autonomic neurorehabilitation (n = 5; independent samples t-test; t = 12.71; p-value = 2.78e-06).

**Supplementary Fig. 10.**
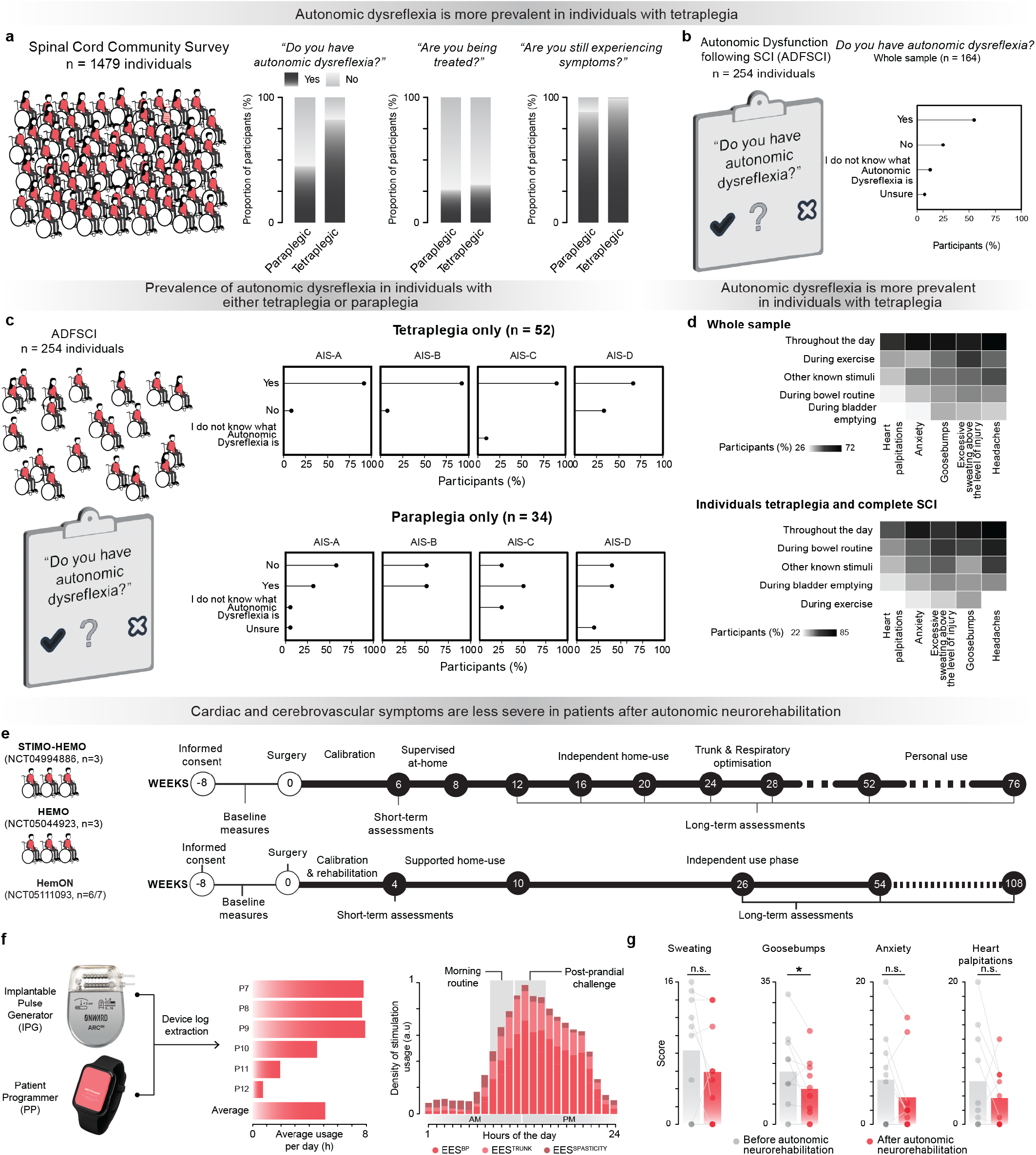
Population-level data of self-reported experiences of autonomic dysreflexia symptoms and clinical validation of autonomic neurorehabilitation. **a**, The prevalence of autonomic dysreflexia and management efficacy in people with SCI (n= 1479) acquired with the Spinal Cord Injury Community Survey (SCICS). **b**, Percentage of individuals with SCI experiencing autonomic dysreflexia scored in the ADFSCI (n=107). **c**, Percentage of autonomic dysreflexia in individuals with spinal cord injuries split between tetraplegia (top) (n = 52) and paraplegia (bottom) (n = 34) acquired with the SCICS. **d**, Percentage of individuals experiencing each symptom described in the ADFSCI autonomic dysreflexia section split between all participants (top) and tetraplegic individuals with complete SCI (bottom). **e**, Timeline of the two clinical trials conducted in Lausanne, Switzerland and in Calgary, Canada. **f**, Bar plots reporting the average daily usage of the system per participant (*left*), and the usage of the system throughout the hours of the day (*right*) for the 8 participants. **g**, Bar plots reporting the ADFSCI autonomic dysreflexia score for each symptom before implantation and at the latest timepoint of home use of the system to regulate blood pressure (n = 10, paired samples one-tailed t-test; t = 1.44, p-value = 0.091; t = 2.19 p-value = 0.028; t = 1.70, p-value = 0.06, t = 1.16, p-value = 0.14).

